# Unbiased genome-scale identification of *cis*-regulatory modules in the human genome by GRAMc

**DOI:** 10.1101/468405

**Authors:** Catherine L. Guay, Jongmin Nam

**Affiliations:** Center for Computational & Integrative Biology, The State University of New Jersey, Camden, NJ 08102, USA; Department of Biology, Rutgers, The State University of New Jersey, Camden, NJ 08102, USA

## Abstract

Although significant advances have been made toward functionally identifying human regulatory elements, existing genome-scale reporter methods preferentially detect either enhancers or promoters. Here we develop GRAMc, a highly reproducible unbiased Genome-scale Reporter Assay Method for *cis*-regulatory modules (CRMs). GRAMc combines the versatility of traditional reporter constructs and the scalability of DNA barcode reporters, and unites the complementary advantages of several currently available high-throughput reporter assays. We demonstrate that GRAMc can reliably measure *cis*-regulatory activity of nearly 90% of the human genome in 200 million HepG2 cells with randomly fragmented ~800bp inserts. By using the GRAMc-identified CRMs, we show that CRMs identified in one cell type are useful for predicting gene regulatory programs not only within that cell type but also between cell types or conditions separated in time and space. In addition, the GRAMc-identified CRMs support the hypothesis that SINE/Alu elements are rich sources of regulatory evolution. Finally, the observation that the majority of experimentally identified regulatory elements do not overlap with computationally predicted elements underscores the necessity of an efficient and unbiased genome-scale reporter assay.

## Introduction

Cis-regulatory modules (CRMs) such as enhancers and promoters are the major functional elements in the genome. It has been estimated that hundreds of thousands of CRMs are scattered across the human genome ^1–3^. Because CRMs regulate when, where, and to what level genes are expressed, CRMs are involved in nearly every biological process. Individual CRMs directly interact with multiple transcription factors, and mutliple CRMs function in combination to mediate gene regulatory activities ^4–6^. Despite the estimated prevalence and biological importance of CRMs, comprehensive experimental identification of these elements remains a challenge.

The standard reporter assay to identify CRMs is to clone a candidate CRM upstream of a basal promoter and a reporter gene (e.g., green fluorescent protein) and to examine its ability to drive reporter gene expression ^7–9^. The same reporter construct can be used to monitor how a CRM responds to gene perturbations ^10^ and to mutations in transcription factor binding sites ^11–15^. Activities of CRMs measured by reporter assays can guide *in vivo* functional analysis of CRMs ^16–18^ and the effect of genetic variations in CRMs ^14,19^. However, conventional one-by-one reporter assays are not suitable for analyzing the millions of potential CRMs contained in the genome.

To meet the challenge of scalability, several high-throughput reporter assays have been developed. Arnold and colleagues have developed STARR-seq ^20^, a genome-scale enhancer screening method to massively discover self-transcribing active regulatory DNAs. STARR-seq and related methods have been successfully applied to screen enhancers in the fly ^20^ and human genomes ^21,22^. Other groups have developed methods that take advantage of the virtually unlimited diversity of short (<30bp) DNA barcodes as reporters ^23–27^. The short and uniform length of barcode minimizes reporter specific variations in RNA stability and PCR amplification biases, and even if there is any bias, use of multiple barcode reporters for the same insert dilutes the effect of reporter specific variations. Recently, DNA barcodes have been applied to a genome-scale screening of autonomous promoters by cloning random human DNA fragments in a promoter-less vector containing barcoded DNA reporters ^28^. It is important to note that currently available genome-scale reporter assays preferentially identify either enhancers or promoters. Several medium-scale methods that use DNA barcode reporters have effectively tested both promoters and enhancers by employing the standard reporter construct design, in which a CRM is placed upstream of a basal promoter and a barcode reporter is downstream ^26,29^. There are other medium-scale methods that rely on the enrichment of active CRMs by fluorescence activated cell sorting ^30,31^. The cell sorting-based methods have the added benefit of massive identification of cell-type specific CRMs from a population of hetertogeneous cells.

Alternatively, CRMs can be predicted based on chromatin signatures that tend to be associated with CRMs ^3^. Experimental validation of predicted CRMs has often focused on testing the accuracy of select groups of candidates that possess multiple signatures and/or trans-species sequence conservation. Depending on how the test CRMs were selected and validated, 12% - 80% of the tested candidates were found to be CRMs in reporter assays ^32–38^ (**Supplementary Table 1**). Although these predictions have considerably accelerated CRM identification, a large number of experimentally discovered CRMs were not associated with these genomic signatures ^21,39^, indicating that currently available prediction tools are not accurate enough to replace reporter assays.

An ideal genome-scale reporter assay method should combine capabilities of currently available methods and overcome their limitations. First, it should be effective for both enhancers and promoters as in the case of standard reporter assays. Second, it should be able to accommodate long DNA inserts, enabling screening of complete CRMs rather than partial CRMs. Third, it should be able to easily control the genomic coverage and the number of DNA barcodes in a library. Excessive genomic coverage and DNA barcodes will increase experimental cost, while insufficient genomic coverage and DNA barcodes will result in less reliable data. Fourth, it should generate reproducible data with comparable or less input materials than currently available methods.

Here, we present a new Genome-scale Reporter Assay Method for CRMs (GRAMc) that combines the versatility of traditional reporter constructs and the scalability of DNA barcode reporters. Using the GRAMc protocol, we have generated a library of reporter constructs that covers the human genome ~4 times (4x coverage) with ≥15 million (M) randomly fragmented ~800bp inserts. The total number of unique DNA barcode reporters in this library is approximately 200M. This library was tested in the human liver carcinoma cell line, HepG2, and identified 41,216 nonoverlapping CRMs that are active at least 5 times higher than the background activity. These CRMs possess several expected features of CRMs. We also demonstrate the utility of CRMs for gene regulatory network analysis and evolutionary genomics.

## Results

### GRAMc library building

The GRAMc library is generated by the following procedure (**Fig. 1**). First, random genomic DNA fragments are size-selected, adapter-ligated, and serially diluted to reach an intended genomic coverage (**Fig. 1a**). To improve the accuracy of adapter ligation, we use a fused adapter to form circular ligation products that can resist Exonuclease I/III treatment against linear DNAs including non-ligated DNAs and linear concatenates (**Supplementary Fig. 1**). After exonuclease treatment, circular ligation products are linearized by RNase HII that cuts ribonucleotide sites (UU/AA) within the fused adapter. Linearized ligates are then serially diluted and PCR amplified using adapter-specific primers. A dilution of intended genomic coverage is identified by counting the presence or absence of 11 randomly chosen genomic regions by QPCR. Note that for a dilution that contains ~4M randomly sampled genomic DNA fragments of 800bp-long (an average of 1x genomic coverage), the expected presence rate of target regions is 0.6. A dilution of 5x or any desired genomic coverage is assembled with two common pieces of DNAs to form a library of linear DNA products that contain genomic test fragments, a basal promoter, a GFP ORF ^40^, and vector backbone (**Supplementary Fig. 2**). The vector system uses a pan-bilaterian Super Core Promoter 1 (SCP) ^41^. Second, the resulting genomic DNA library is barcoded with an excess number of random 25mers (N25) by PCR with a pair of common primers that can amplify the entire library including the vector backbone (**Fig. 1b**). One of the common primers, primer_R, contains random N25 in the middle and a core-poly adenylation signal (polyA) ^42^. The barcoded library is self-ligated, Exonuclease I/III treated and electroporated into *E. coli* for library amplification and plasmid extraction. A small fraction (e.g., 1/1,000th) of unrecovered transformants is used to measure the colony forming unit (cfu), and the remainder is used for library amplification in liquid culture and subsequent plasmid extraction. Because the PCR-mediated barcoding introduces an excess of barcodes, virtually all individual transformants contain unique barcodes. For example, barcodes present in transformants used for colony counting were not identified in the final library. The number of unique barcode reporters in a GRAMc library can be controlled by the scale of electroporation. In our protocol, 4 - 10ng of circular ligation products with ~800bp inserts consistently generated ~40M cfu, which is comparable to the advertised efficiency of commercially available competent cells. Note that the genomic coverage of the library that was determined in the first step will be maintained, as long as the number of unique barcodes harvested is much larger than the number of unique inserts. Purified plasmids are used for library characterization. Library characterization includes identification of genomic inserts and pairs of inserts and barcode reporters by Illumina paired-end sequencing (See **Methods** and **Supplementary Fig. 3**).

**Figure 1.**
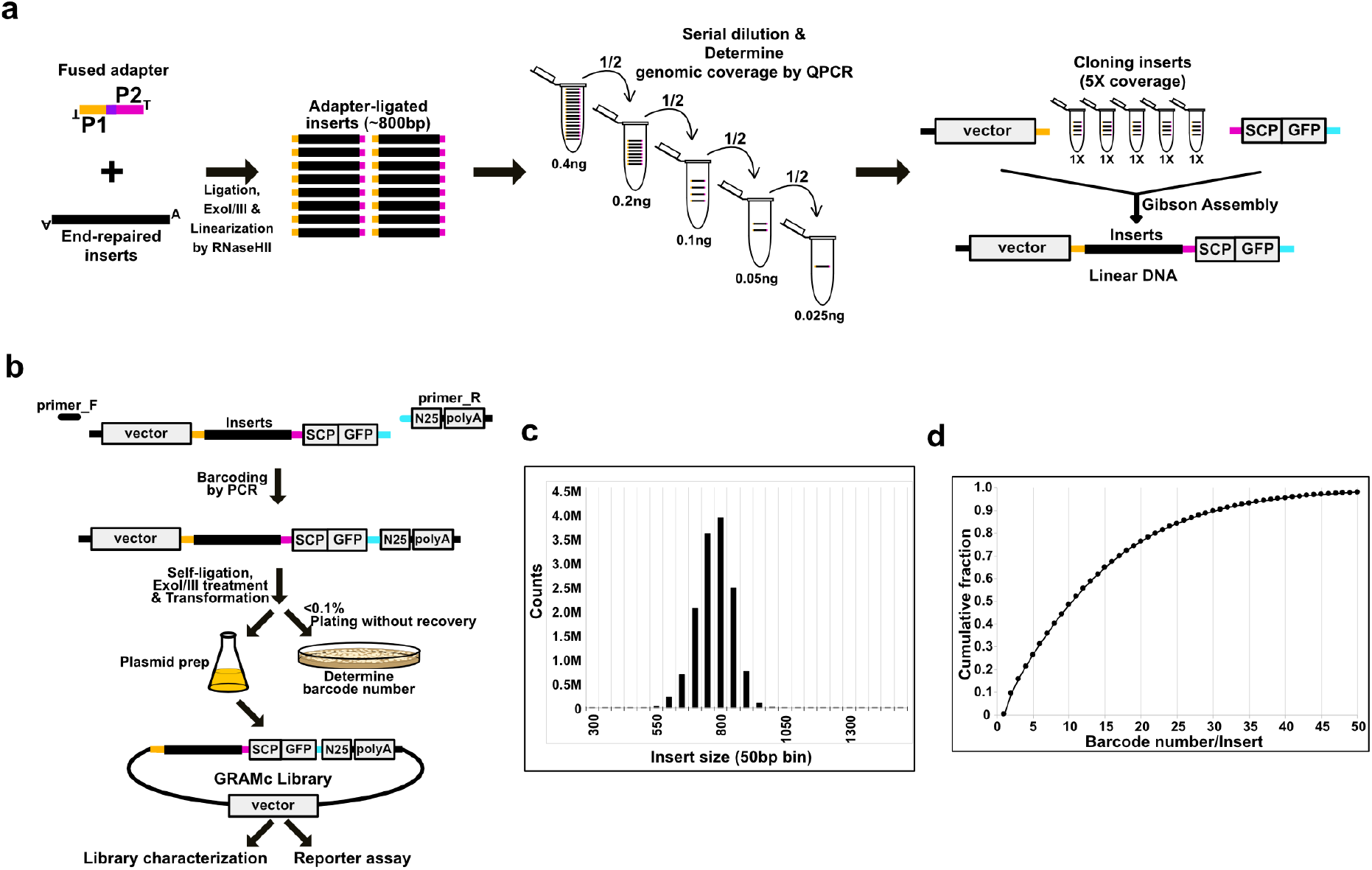
GRAMc library building. (**a**) Controlling genomic coverage of the library. Size-selected and end-repaired random genomic DNA fragments are circularized by ligation with a fused adapter. Linear DNAs are removed by exonuclease treatment followed by RNaseHII digestion to linearize ligation product and dice adapter-concatemers. The purple box indicates two ribonucleotides for RNaseHII cut. Adapter-ligated products are then serially diluted to determine the genomic coverage of each dilution by QPCR. A dilution of intended coverage is Gibson assembled with a SCP-GFP cassette and the vector backbone to form barcode-less, linear constructs. (**b**) Controlling barcode numbers of the library. Random 25bp (N25) barcodes and a core-poly adenylation signal are added to the library of linear constructs by PCR. Barcoded constructs are self-ligated and linear DNAs are removed by exonucleases I/III. A small fraction of ligates is transformed to determine the scale of transformation. To avoid inflation of colony counts due to cell division, transformants for counting colonies should be immediately plated without rescuing. A desired amount of ligates are transformed to produce a GRAMc library with the intended number of barcodes. Plasmids extracted from liquid media are used for library characterization and reporter assay. Inserts and associated barcodes are identified by Illumina paired-end sequencing. (**c**) Size distribution of inserts in the human GRAMc library. (**d**) Cumulative distribution of barcode numbers per insert in the human GRAMc library.

Using this method, a human GRAMc library of ~800bp-long inserts was generated. The intended numbers of unique genomic DNA inserts and unique barcodes in this library were 20M (5x genomic coverage) and 200M (10 barcodes/insert), respectively. After analyzing 479.1M pairs of sequences mapped to the hg38 assembly (out of 519M paired-end reads), 15.1M genomic regions were identified. The total number of unique barcodes that are associated with these genomic regions was 191M. This library covers 93.4% of the human genome at least once (**Supplementary Table 2**). Although obtaining more sequencing reads would improve these numbers, these numbers are already close to the intended yield of inserts and barcodes in the library. Note that, of the detected 15.1M genomic regions, 13.2M inserts were unique in sequences (<95% sequence identity with other genomic regions). In addition, the genomic distribution of unique inserts was more or less uniform (**Fig. 2c**). When we checked the lengths of unique inserts (**Fig. 1c**), 71% of the inserts were within 750 - 850bp range, suggesting that our size selection was effective. In the case of the number of barcodes per insert (**Fig. 1d**), although the barcode numbers of the majority of inserts deviated significantly from the expected number of 10, 99% and 55% of unique inserts were connected to ≥2 barcodes and ≥10 barcodes, respectively. Therefore, barcode-specific effects on reporter expression will be insignificant in the GRAMc library.

**Figure 2.**
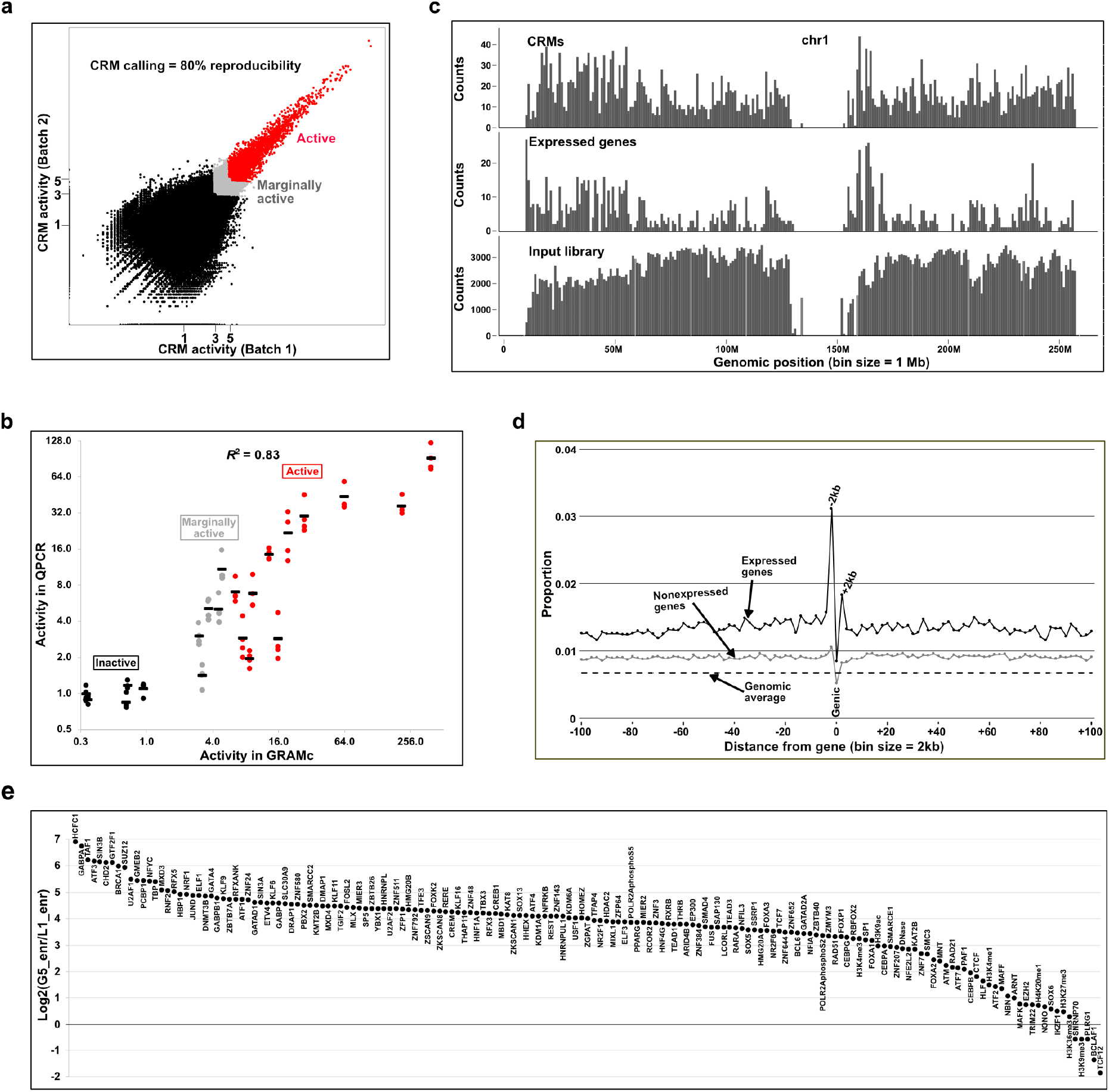
Reproducibility and accuracy of GRAMc. (**a**) Reproducibility of GRAMc results. The human GRAMc library was tested in two batches of 200M HepG2 cells. CRM activities are double-normalized to the copy numbers of input plasmids and background activity (bg). Inserts that drove reporter expression ≥5xbg in one batch and ≥4.5xbg in another are considered CRMs (red dots), and the CRM calling was 80% reproducible. Inserts that did not meet the cutoff but are still ≥3xbg in one batch and ≥2.7xbg in another are called marginally active (grey dots) with a lower reproducibility of 62%. (**b**) Validation of GRAMc results by individual reporter assay. A set of 11 CRMs (red dots), 5 marginally active inserts (grey dots) and 5 inactive inserts (black dots) were tested in 4 batches of individual reporter assays by QPCR. Averaged activities (black bar) from 4 batches of individual reporter assays were compared to the GRAMc data (*R*^2^ = 0.83). (**c**) Correlated genomic distributions of CRMs (top) and expressed genes (middle) on chromosome 1. Genomic distribution of the input library is shown at the bottom. Inserts from centromeres are removed. (**d**) Enrichment of CRMs in 2kb windows up to 100kb flanking regions of expressed genes (black dots) and nonexpressed genes (gray dots). Genomic average is shown as a dashed line. Genic region is at position 0 and includes both exons and introns. Upstream of genes are on the left half and downstream on the right half. (**e**) Relative enrichment of ENCODE chromatin annotations in CRMs (G5, greater than 5xbg) *vs.* inactive inserts (L1, lower than 1xbg). ENCODE annotations are ordered based on their relative enrichment.

### Application of GRAMc in HepG2 cells

The GRAMc library was tested in two batches of 200M HepG2 cells. As a comparison, previous genome-scale enhancer screenings used 300M LNCaP cells ^21^ and 800M HeLa cells ^22^, and a genome-scale promoter screening used 100M K562 cells ^28^. Following transfection of the GRAMc library into cells, total RNAs were extracted, reverse transcribed, and expressed barcodes were PCR amplified. To avoid losing reporter transcripts during secondary enrichment of mRNA ^22^ or reporter transcripts ^35^, we use total RNAs and GRAMc-specific oligomer for reverse transcription. Expressed barcodes were amplified by PCR, and expression levels of reporters were measured by Illumina sequencing. A schematic of processing RNAs into sequencing libraries, along with the associated quality control steps is available in **Supplementary Fig. 4**. Reporter expressions were double-normalized to the relative copy number of inserts in the input GRAMc library and background activities, which is the average activity of the middle 30% of rank ordered reporter expressions ^26^. The background activity measured in this way has been very similar to the leaky activities of known inactive fragments in sea urchin embryos ^26,27^.

Approximately 200M reads from each batch of expressed barcodes were obtained, and 78 - 79% of barcodes matched to barcodes with associated genomic regions. To account for copy number variations, approximately 450M barcode reads were obtained from input plasmids. Because 99% of inserts are driving ≥2 barcodes, read numbers of multiple barcodes for the same insert were combined. Approximately 7.5M inserts with ≥10 reads from input plasmids were used for data analysis. A total of 50,993 inserts from 41,216 non-overlapping genomic regions displayed activities of ≥5-fold higher than the background (bg) activity (≥5xbg) in one batch and ≥4.5xbg in another batch (red dots in **Fig. 2a**). Our replicate GRAMc data showed Pearson’s correlation coefficient (*r*) of 0.95, and the probability of a CRM in one batch being called a CRM in another batch was 0.80 (80% reproducibility of calling CRMs). When the cutoff was lowered to 3-fold of the background (grey and red dots, ≥3xbg in one batch and ≥2.7xbg in another batch), the number of active regions increased to 150,011 (62% reproducibility of calling CRMs). The list of genomic coordinates of inserts and their normalized activities is available as **Supplementary File 1**.

To validate the accuracy of GRAMc, we randomly selected 11 CRMs (≥5xbg, red dots in **Fig. 2a**), 5 marginally active fragments (3-5xbg, grey dots in **Fig. 2a**), and 5 inactive fragments (≤1xbg, black dots in **Fig. 2a**) and individually tested their regulatory activities with a one-by-one reporter assay (**Fig. 2b**). Levels of GFP transcripts relative to copies of transfected DNA were measured by QPCR. Reporter expressions were further normalized to the background activity (bg), which is the averaged level of the 5 inactive reporter constructs. Average levels of 4 independent assays were shown in black bars for individual inserts. Of the 11 CRMs tested 8 inserts were ≥5xbg, while 2 inserts and 1 insert were respectively 2.8xbg and 1.9xbg. This result is comparable to the 80% reproducibility of calling CRMs in GRAMc (**Fig. 2a**). In the case of the 5 marginally active inserts, 1 insert was 10xbg, 3 inserts were within the expected range of 3 - 5xbg, and 1 insert was 1.4xbg. Overall, *cis*-regulatory activities measured by GRAMc were reproducible in independent assays (*R*^2^ = 0.83). The list of inserts selected for individual reporter assays is available in **Supplementary Table 3**. These results indicate that GRAMc is a reliable and efficient tool to discover CRMs at genome-scale.

### GRAMc-identified CRMs possess expected features of CRMs

Since GRAMc is based on the standard configuration of reporter constructs, GRAMc-identified CRMs should possess known features of CRMs that have been identified by traditional reporter assays. Firstly, CRMs should primarily be located near expressed genes in HepG2. When we compared the genomic locations of expressed genes in HepG2, CRMs and the input library, expressed genes and CRMs had similar patterns while the input library was approximately uniformly distributed (**Fig. 2c and Supplementary Fig. 5**).

Secondly, CRMs are known to be enriched 5’-proximal to genes (promoters), however the majority are located outside of the proximal regions (distal enhancers) ^26^. When we computed the proportions of CRMs for the number of inserts tested within sliding 2kb windows upstream or downstream of expressed genes, the 5’-proximal 2kb regions showed the highest enrichment (0.03) (**Fig. 2d**). The 3’-proximal 2kb regions showed the second highest peaks, while genic regions are slightly depleted of CRMs. Despite these regional variations, CRMs are consistently enriched around expressed genes within at least 100kb region in each direction compared to the genomic average of 0.007. A similar pattern was also observed near unexpressed genes, but the degree of enrichment was lower than near expressed genes. These results indicate that GRAMc can efficiently identify both proximal promoters and distal enhancers.

Thirdly, CRMs are expected to be associated with bindings of transcription factors and other proteins that positively impact CRM function. When we computed the relative enrichment (total base pairs shared relative to random expectations) of narrow peaks from 167 data sets of ENCODE ChIP-seq or DNase-seq from HepG2 in CRMs *vs.* inactive fragments (**Fig. 2e**), 153 data showed ≥2-fold enrichment in CRMs *vs.* inactive regions. These include general transcriptional factors (e.g., GTF2F1, TAF1 and TBP), a transcriptional coactivator (P300) and histone modification enzymes (e.g., H3K4me3 and H3K9ac). ChIP-seq peaks that were not enriched or were even depleted in CRMs include transcription factors (TCF12 and BCLAF1), spliceosome components (PLRG1 and SNRNP70), and histone methylases (H3K27me3, H3K36me3 and H3K9me3). Interestingly, despite the overall enrichment, only 32% of GRAMc-identified CRMs overlapped with the 153 ENCODE data with ≥2-fold enrichment in CRMs, and 58% of CRMs did not overlap with any ENCODE data used in this analysis. Although we expect that obtaining ChIP-seq data for more transcription factors would increase the overlap, it is also possible that reporter assays can detect CRMs that are not active in the genome due to chromatin silencing or CRMs that can evade detection by ChIP-seq and chromatin marks. The list of ENCODE data used in this analysis is available in **Supplementary File 2**.

### Motif enrichment explains differential activities of ChromHMM predicted enhancers

Earlier studies have shown that, although CRM predictions based on chromatin marks are enriched in functionally validated CRMs, the majority of predicted CRMs did not drive significant expression in reporter assays ^21,22,28^. To test whether this is also the case in GRAMc, we checked *cis*-regulatory activities of GRAMc-tested inserts that overlap ≥90% with ChromHMM-predicted strong enhancers in HepG2 ^43^ (**Fig. 3a**). The result shows that predicted enhancers were 3.3-fold enriched in the cohorts of inserts that were ≥5xbg (G5 set) and ≥3xbg and <5xbg (G3L5 set). Even the cohort of inserts with ≥2xbg and <3xbg (G2L3 set) showed 2.2-fold enrichment of predicted enhancers. However, the majority of inserts (80%) that overlap with predicted enhancers did not drive significant reporter expressions, and belong to G1L2 or L1 sets. We hypothesized that if the predicted enhancers were true enhancers, one would expect enrichment of transcription factor binding site (TFBS) motifs. Note that we focused on predicted strong enhancers, because promoters are inherently enriched with TFBS motifs and predicted weak enhancers may increase ambiguity.

**Figure 3.**
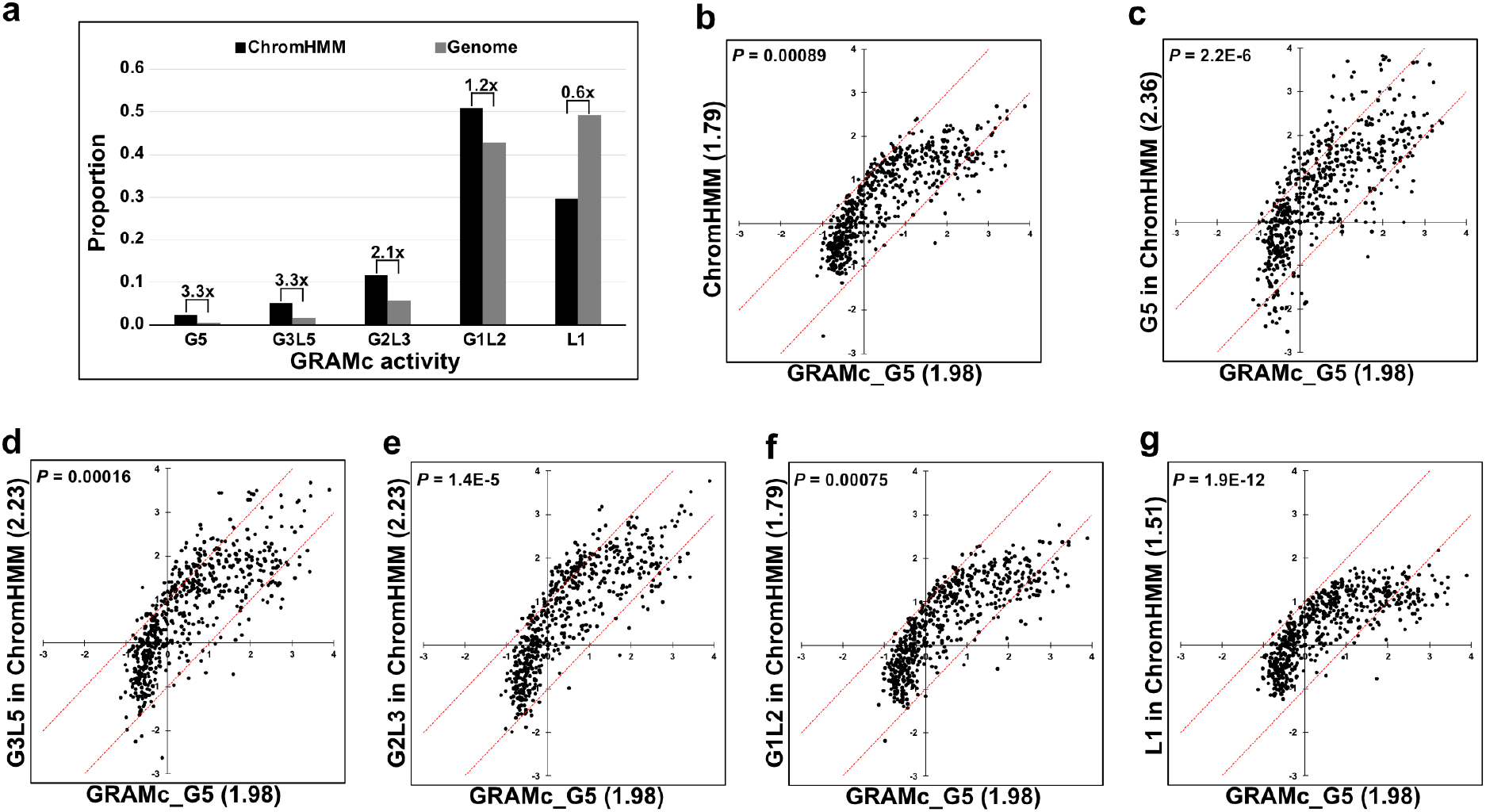
Cis-regulatory activity and TFBS motif enrichment in ChromHMM-predicted strong enhancers. (**a**) Relative enrichment of ChromHMM-predicted strong enhancers (black bars) *vs.* genomic averages (gray bars) in different cohorts of GRAMc activity. Fold enrichment of predicted enhancers *vs.* genomic average is shown above each pair of bars. Cohorts were grouped based on the averaged activity in two batches of GRAMc data: G5, equal or greater than 5xbg; G3L5, equal or greater than 3xbg and lower than 5xbg; G2L3, equal or greater than 2xbg and lower than 3xbg; G1L2, equal or greater than 1xbg and lower than 2xbg; L1, lower than 1xbg. (**b** - **e**) Relative motif enrichments (Log2 scale) in predicted enhancers with progressively weaker GRAMc activities *vs.* GRAMc-identified CRMs (G5). The average enrichment value of each cohort before log transformation is shown within parenthesis, and the *P*-value for each comparison was computed by a Paired *t*-test (two-tailed). Each dot represents a TFBS motif and red lines indicate 2-fold differences between the two data sets.

To test the hypothesis, we compared enrichments of 601 HOCOMOCO_v10 HUMAN motifs ^44^ within predicted enhancer regions and GRAMc-identified CRMs using inactive fragments as negative control. Overall, when we compared the average enrichment values (**Fig. 3b**), GRAMc-identified CRMs showed a higher enrichment of motifs than did the predicted enhancer regions (*P* = 0.00089, Paired *t*-test). Interestingly, the predicted enhancers in G5, G3L5, and G2L3 sets showed higher average enrichment values than that of the GRAMc-identified CRMs (**Fig. 3c-e**), suggesting the possibility that GRAMc data and chromatin marks may complement each other. However, enrichment of motifs gradually decreased in predicted enhancers in G1L2 and L1 sets (**Fig. 3f & g**). Given their inability to drive significant reporter expressions and weak motif enrichment, the predicted enhancers that belong to G1L2 and L1 sets are unlikely to be true enhancers. However, this does not rule out the possibilities that chromatin marks may indicate a neighborhood of enhancers rather than the exact location, and that predicted enhancers may possess other types of *cis*-regulatory activities that cannot be measured in reporter assays. The list of sequences used for motif enrichment analysis is available in **Supplementary File 3**.

Recently, it was shown that activation of the interferon pathway upon DNA transfection results in erroneous identification of interferon-responsive enhancers ^22^, and such an artefact can reduce overlap between GRAMc-identified CRMs and ChromHMM predictions. However, consistent with the original discovery that HepG2 cells do not activate the pathway, motifs for interferon-stimulated transcription factors including IRF1 - 9 and hMX1 were not enriched in GRAMc-identified CRMs (**Supplementary File 4**).

### Enriched motifs in CRMs predict gene regulatory interactions within and between cell types

The pattern of reporter expressions measured by small reporter constructs are direct readouts of the trans-regulatory environment in host cells. Because the DNA sequences of CRMs contain binding sites for transcription factors, computational motif analysis has often been used to infer gene regulatory programs e.g., ^45,46–49^. Based on 601 HOCOMOCO_v10 HUMAN motifs ^44^ computationally predicted in the CRMs and in inactive fragments (negative controls) by FIMO ^50,51^, we computed Abundance (the proportion of motif-positive CRMs or inactive fragments) and Relative enrichment of motifs (relative abundance of a motif in CRMs *vs.* inactive fragments) (**Fig. 4a**) (**Supplementary File 4**). We detected that 176 out of 601 motifs were ≥2-fold enriched in CRMs compared to inactive fragments. While the majority of enriched motifs (65%) were for expressed (Fragments Per Kilobase of transcript per Million mapped reads, FPKM ≥1) transcription factors, interestingly, the remainder were for nonexpressed or very lowly expressed (FPKM < 1) transcription factors ^3^.

**Figure 4.**
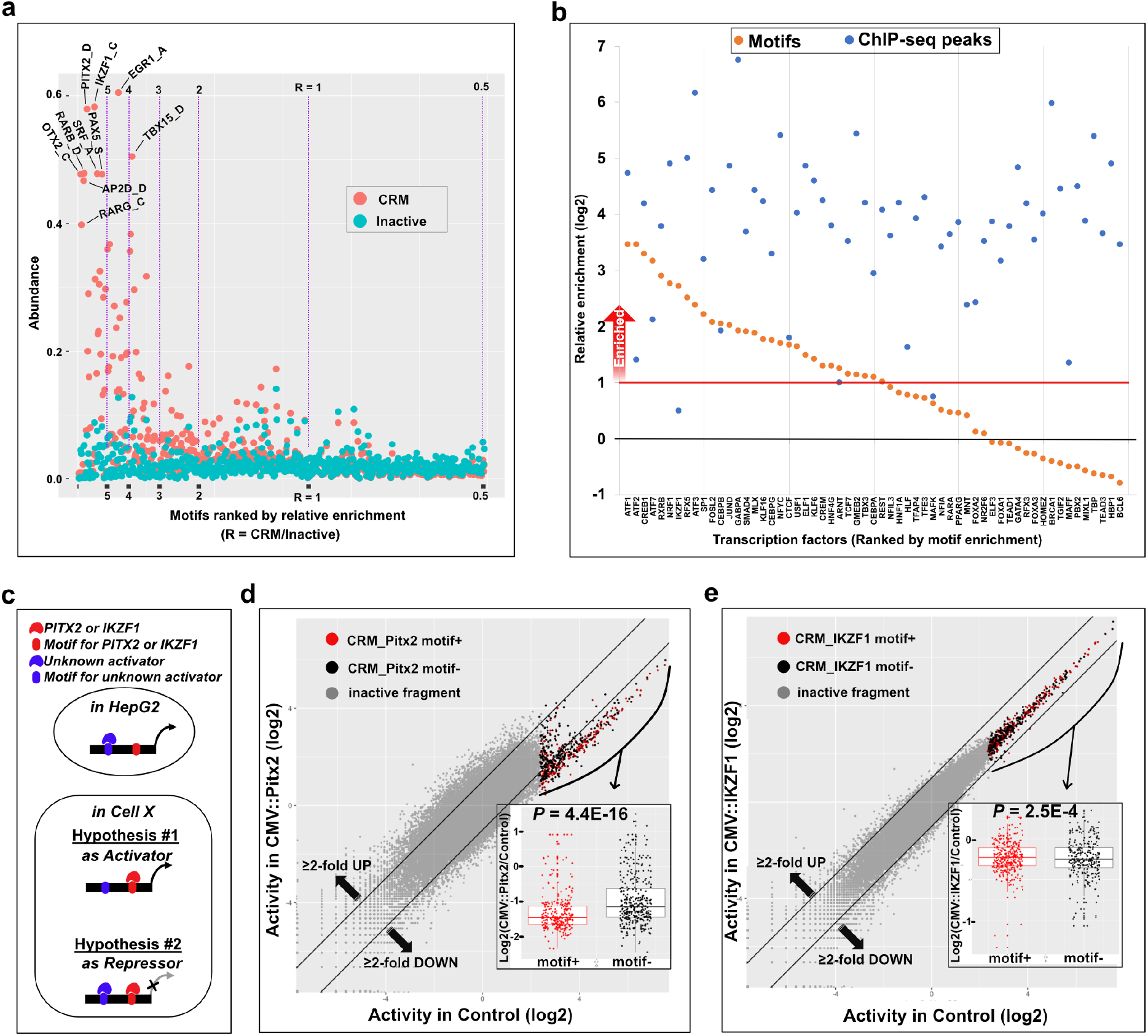
CRM-driven prediction of gene regulatory programs. (**a**) Abundance and enrichment of TFBS motifs in CRMs. Abundance is the proportion of CRMs (the G5 set, orange dots) or inactive sets (the L1 set, cyan dots) that contain a given TFBS motif, and the relative enrichment is the ratio of motif enrichments between the G5 set and the L1 set. Purple vertical lines indicate borders for the relative enrichment of motifs. Several highly enriched and abundant motifs are labeled. (**b**) Comparison of the enrichments of predicted TFBS motifs (yellow dots) and ENCODE ChIP-seq annotations (blue dots) in the G5 set. (**c**) Two alternative hypotheses on the role of PITX2 or IKZF1 on HepG2-CRMs in other cells (Cell X). (**d** & **e**) Testing a hypothesis on the enriched TFBS motifs for non-expressed transcription factors in HepG2 by ectopic expression of human *pitx2* (**d**) and human *ikzf1* (**e**) *vs.* CMV::gfp control. Inserts that belong to the G5 set are shown in red dots (motif+) or in black dots (motif-). Two black diagonal lines indicate 2-fold differences between the perturbed set vs. the control set. Inset boxplots show the different between motif+ vs. motif-inserts with *P* values by Two-Sample *t*-test.

We reasoned that enriched motifs for expressed transcription factors predict positive regulators for the CRMs identified in HepG2. If a motif-based prediction of positive regulators is correct, we would expect enrichment of ChIP-seq peaks for the same transcription factor. To check the accuracy of the prediction, we compared our motif analysis result with ENCODE ChIP-seq data from HepG2 cells ^3^. Regardless of motif enrichment, a total of 58 transcription factors were shared between the two datasets. Of the 31 transcription factors with ≥2-fold enrichment of motifs in CRMs (orange dots above the red horizontal line in **Fig.4b**), all but one transcription factor (IKZF1) had ChIP-seq peaks enriched in CRMs (blue dots in **Fig.4b**). Interestingly, the transcript of *ikzf1* was not detected in HepG2 by RNA-seq ^3^, suggesting that the ChIP-seq peaks may not be true IKZF1 binding regions. These results indicate that the motif-based prediction of positive regulators has a very low false positive rate. In the case of 27 transcription factors with <2-fold enrichment of motifs in CRMs (orange dots below the red horizontal line in **Fig.4b**), 26 factors had ChIP-seq peaks enriched in CRMs (blue dots in **Fig.4b**). Although a more detailed analysis is necessary to explain the latter observation, in the worst-case scenario, the false negative rate of motif-based prediction of positive regulators is estimated to be ~0.5.

Enrichment of motifs for nonexpressed transcription factors led us to hypothesize that they control the HepG2-CRMs either as an activator or as a repressor in other cell types or conditions (**Fig. 4c**). We tested this hypothesis by ectopic expression of candidate transcription factors in HepG2. We examined two transcription factor genes, *pitx2* (a homeobox gene) and *ikzf1* (an ikaros homolog). In mice, *pitx2* is expressed in and is required for hematopoietic function of the fetal liver ^52^, and suppression of *pitx2* expression in induced hepatic stem cells promotes differentiation of hepatocytes or cholangiocytes ^53^. Similarly, *ikzf1* is a key regulator of hematopoietic development ^54^ and expressed in the fetal liver ^55^, although its function in hepatic development is not known. We co-transfected plasmids that can constitutively express mRNAs of *pitx2* (CMV::pitx2) or *ikzf1* (CMV::ikzf1) with a set of randomly selected ~80,000 GRAMc reporter constructs from the full GRAMc library. As a control experiment, we co-transfected plasmids that can constitutively express GFP mRNAs (CMV::gfp) with the same set of reporter constructs. Replicate experiments of all three experiments were highly reproducible (Pearson’s *r* ≥ 0.99) (**Supplementary Fig. 6**). Ectopic expression of *pitx2* in HepG2 down-regulated the majority of CRMs by ≥2-fold and this down-regulation was more pronounced in pitx2 motif-positive CRMs (Two-Sample *t*-test, *P* = 4.4E-16) (**Fig. 4d**). In the case of *ikzf1*, only 9 CRMs were downregulated by ≥2-fold and 6 of the 9 down-regulated CRMs were positive for IKZF1 motif (Two-Sample *t*-test, *P* = 2.5E-4) (**Fig. 4e**). These results suggest that *pitx2* (and *ikzf1* to a minor degree) keeps HepG2-CRMs repressed in the fetal liver, and clearance of *pitx2* is critical for the activation of HepG2-CRMs and gene expression in the adult liver. Although this model needs further validation, our results indicate that CRMs are not only useful to predict regulatory programs in the host cell, but also to predict regulatory interactions between cells separated in time and space.

### SINE/Alu elements are enriched in CRMs

Early models for eukaryotic gene regulation proposed repeat elements as a key player in gene expression control ^56,57^. These predictions were later supported by multiple examples of Alu and ERV elements contributing to gene regulation and its evolution ^58^. Recently, genomic surveys of chromatin signatures have shown that SINE/Alu elements are enriched in putative CRMs ^59,60^. However, genome-scale reporter assays for enhancers ^22^ or promoters ^28^ have detected enrichment of LTR/ERV1 and LTR/ERVL-MaLR in CRMs but not SINE/Alu. To check whether this is also the case in GRAMc-identified CRMs, we have compared our data with annotated repeat elements in the human genome ^61^. We have detected three families of repeat elements, Satellite/telomere, SINE/Alu and LTR/ERV1, that are enriched ≥2-fold in CRMs (G5 set in **Fig. 5a**), but LTR/ERVL-MaLR was not enriched in CRMs. The three elements were also enriched in marginally active G3L4 and G4L5 sets to lesser degrees. Interestingly, alpha-satellites were depleted ~8-fold in CRMs, suggesting their repressive function or incompatibility with CRMs in HepG2. However, we did not detect depletion of Retroposon/SVA elements that were predicted to be transcriptional repressors in liver ^62^.

**Figure 5.**
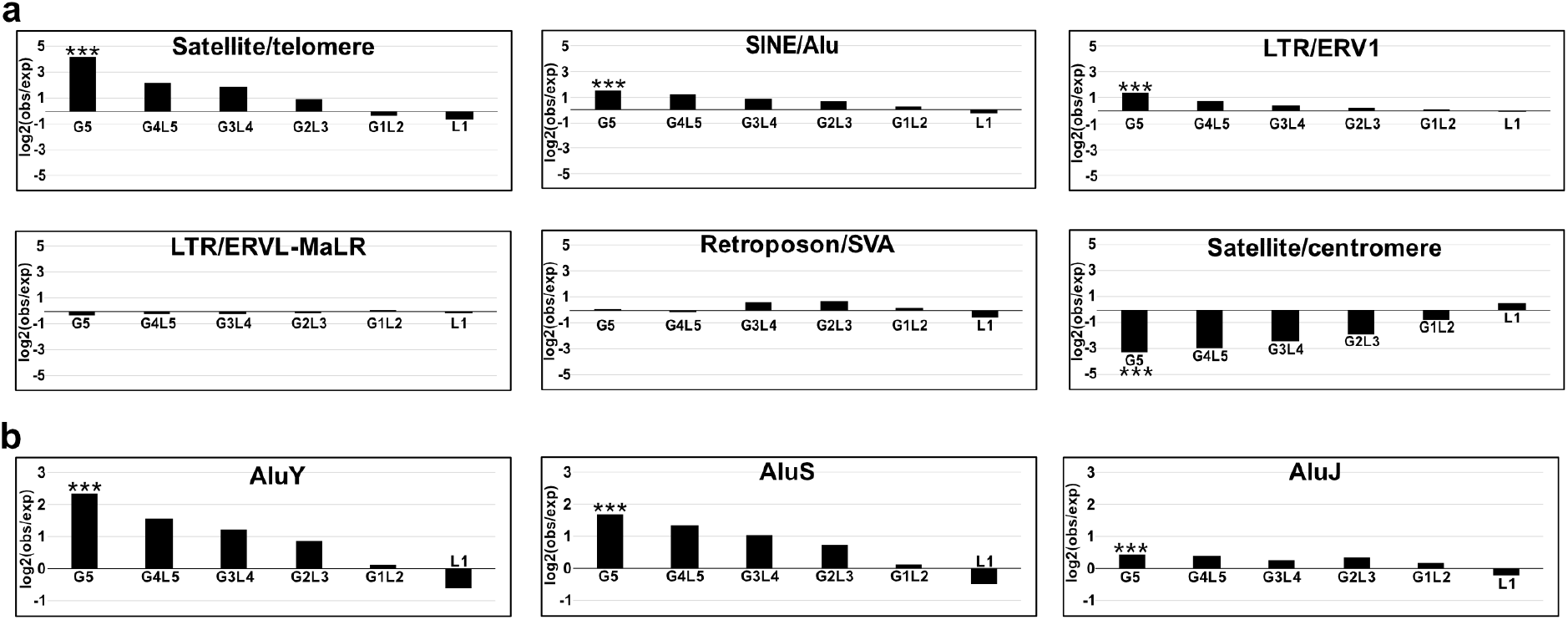
Enrichment of repeat elements in GRAMc data. Inserts were classified by their averaged activities in two batches of GRAMc data as in the case of Figure 3. (**a**) Representative families of repeat elements in GRAMc data. Enrichment of repeat elements within genomic regions with differential activities. Genomic regions in the G5 set are considered CRMs. (**b**) Enrichment of three major subfamilies of Alu elements in GRAMc data. The G5 cohorts with statistically significant (*P* < 0.01) enrichment or depletion of repeat families are marked with “***”.

Using the GRAMc-identified CRMs, we also tested a recently proposed model that Alu elements have evolved toward enhancers over time ^59^. This model predicts that the enrichment of Alu elements in CRMs should be positively correlated with their age. However, when we checked three major subfamilies of Alu (**Fig. 5b**), the youngest subfamily (AluY) and the intermediate subfamily (AluS) showed ≥3-fold enrichment in CRMs, while the oldest subfamily (AluJ) showed only moderate enrichment (1.3-fold). Because the original study is based on the chromatin annotations in HeLa cells, this discrepancy can be explained by differences in cell types. To check this possibility, we compiled subfamilies of 19 Alu elements that were tested with luciferase assays in HeLa cells ^59^. Consistent with our results, 8/10 AluY or AluS elements were active and only 4/9 AluJ elements were active. Therefore, our results are consistent with an alternative model that Alu elements lose regulatory activity with age.

These results demonstrate that GRAMc data can be useful for testing multiple evolutionary genomics hypotheses, and that it can lead to different conclusions compared to the data generated by earlier genome-scale reporter assays or chromatin annotations. Note it is possible that the observed discrepancies between GRAMc and earlier reporter assays attributed in large part to different cell types used. Enrichment of the entire list of repeat elements is available in **Supplementary File 5**.

## Discussion

GRAMc combines complementary advantages of multiple existing large-scale reporter assay methods and is free from prior assumptions other than the fundamental assumption that a CRM should be able to drive reporter gene expression by activating a heterologous basal promoter. We have demonstrated that GRAMc can reliably identify both enhancers and promoters from the entire human genome in HepG2 cells.

Comparison of GRAMc data and chromatin marks solidifies the recurring observation that, despite statistically significant enrichments, the majority of CRMs do not overlap with available chromatin marks and *vice versa*. Are the majority of GRAMc-identified CRMs an artefactual outcome of mere abundance in TFBS motifs rather than their actual function in the genome? If this is the case, one would expect that CRMs would be equally enriched near genes regardless of their expression. Because GRAMc-identified CRMs are preferentially enriched near expressed genes, we think this is unlikely. An alternative scenario is that overlap or nonoverlap between GRAMc-identified CRMs and chromatin marks may indicate different mechanisms of their activation or function. A key question that arises from this scenario is whether GRAMc-identified CRMs that overlap with positive chromatin marks are more biologically relevant than CRMs that do not overlap with the positive marks. Because the current version of GRAMc is an episomal reporter assay, chromatin-level regulations that are not originated by the CRM in question are likely to be missed. In a previous study ^63^, a few chromatin annotations including SIN3B and BRCA1 were preferentially associated with CRMs that were stronger in episomal assays *vs.* chromosomally integrated reporter assays. Interestingly, the same chromatin annotations were among the highest enriched in GRAMc-identified CRMs (**Fig. 2e**). Availability of a comprehensive set of functional CRM data that does not rely on chromatin annotations should be useful to elucidate this discrepancy and to better utilize available data to test various functional and evolutionary genomics hypotheses ^64^.

Regardless of their functionality in the genome, individual CRMs in reporter constructs interact with a subset of transcription factors that are active in a cell. Therefore, a large number of CRMs in reporter constructs are essentially reading devices of entire gene regulatory programs. Our analysis demonstrated that simple motif analysis could predict a substantial portion of gene regulatory programs with a very low false positive rate. Given that a CRM can contain regulatory information for multiple cell types in its DNA sequence, our test with ectopically expressed *pitx2* and *ikzf1* in HepG2 cells suggested that CRMs can inform gene regulatory programs between cells that are separated by space and/or time. Because nearly 1/3 of enriched motifs HepG2-CRMs were for transcription factors that are not expressed in HepG2 cells, this type of higher-order gene regulatory interactions should be prevalent, especially between lineage-related mutually exclusive cell types/conditions.

After detailed comparison with other methods ^20–22,28,35,63,65^, we realized that there are several common steps in which we have improved yield/efficiency by 2 - 5 folds. These steps include *i*) using proteinase K to reduce viscosity of ligation reaction during library building, which improves recovery of ligation products nearly 2-fold (Method step 1), *ii*) preparing a pure population of circular ligates to achieve a commercially advertised transformation efficiency during library building (Method step 1), *iii*) adding Exonuclease I/III to complement low activity of DNase I for single stranded DNAs to rigorously remove DNAs in RNA samples (Method step 3), and *iv*) using 4mg of total RNAs for reverse transcription without secondary RNA enrichment (Method step 3). These steps are modular and can easily be implemented into other methods. Conversely, the GRAMc vector can be modified to use plasmid origin of replication as basal promoter to boost reporter expression ^22^. While the effect of individual improvements may be small, the collective improvement of the efficiency and accuracy will be critical for application of genome-wide reporter assays to cells that are much harder to transfect than well-established cell lines.

## Acknowledgments

The authors thank Dibyendu Kumar and Brian Gelfend at Waksman Genomics Core for Illumina sequencing; the Caliburn team for maintaining the Caliburn cluster; Thomas Skipper for maintaining the vSMP and Golova clusters; the ENCODE Project for making their unpublished data freely available; Eric Klein for comments on an earlier version of the manuscript. This work was supported by Start-up and Faculty Research and Creative Activities Program from Rutgers and by New Jersey Health Foundation (PC90-17) to JN.

## Author contributions

JN conceived the study. CLG and JN did the experiment. JN did the data analysis. CLG and JN wrote the paper.

## Methods

### 1. GRAMc library construction

#### Fused adapter preparation

GRAMc uses a custom-designed fused adapter to minimize the formation of unwanted concatenates (**Supplementary Fig. 1**). Two complementary hybrid oligomers were synthesized by Integrated DNA Technologies (IDT): p-AD4_F (5’-/p/CTGCTGAATCACTAGTGAATTATTACCCrUrUCAAGACACTACTCTCCAGCAGT-3’) and p-AD4_R (5’-/p/CTGCTGGAGAGTAGTGTCTTGrArAGGGTAATAATTCACTAGTGATTCAGCAGT-3’). Ribonucleotide sites are labeled “rU” and “rA.” A fused adapter was prepared by diluting p-AD4_F and p-AD4_R to 4pmol/uL in 1x T4 DNA ligase buffer (NEB B0202S) followed by annealing at 95C for 2 min, then decreasing the temperature for 160 cycles at a rate of −0.5C/20s cycle. Annealed adapters were aliquoted into 3ul volume and kept in −80C until use.

#### GRAMc vector preparation

The GRAMc vector was constructed by replacing the sea urchin nodal basal promoter with the Super Core Promoter 1 (SCP) ^1^ upstream of the GFP ORF in an existing vector ^2^ based on pGEMT Easy vector (Promega). The GFP ORF is from pGreen Lantern (Gibco BRL) ^3^. The template vector was linearized by AflII/HindIII overnight digestion and amplified in 10 cycle of PCR as two separate cassettes from 20ng of linearized template (**Supplementary Fig. 2**). The SCP-GFP cassette was amplified in a 50uL Q5 High-Fidelity DNA Polymerase reaction (NEB M0491) using primers NJ-95 and NJ-145 and the vector backbone with NJ-146 and NJ-96 using an annealing temperature of 62C and a 2min extension. A sequence of six phosporothioated bases at the 5’ end of the NJ145 and NJ146 prevent loss of primer sites during subsequent Gibson Assembly.

#### Preparation of genomic inserts

We randomly fragmented 20ug of NG16408 genomic DNA (Coriell Institute) in 200uL of water with a Qsonica Q125 at 20% Amperage with 3 cycles of 15s pulses/10s rest. We column cleaned DNA using a Zymo-25 column (Zymo Research), and size selected ~800bp fragments on a 1.2% agarose gel. A portion of the gel purified gDNA was size confirmed on a 2% Agarose E-gel (ThermoFisher G501802). The remaining purified fragments were repaired in a 25uL PreCR reaction (NEB M0309) containing 1X Thermopol Buffer, 100uM dNTPs, 1X NAD+, and 0.5uL of PreCR enzyme for 30 minutes at 37C. PreCR treated fragments were column purified using a Zymo-6 column, and treated with the End Repair/dA Tailing Module (NEB E7370) in a 32.5uL reaction, followed by a 41uL reaction of the TA Ligation Module (NEB E7370) with a 10:1 adapter to insert molar ratio of the annealed AD4 fused adapter. Unligated adapters and genomic inserts were removed with 20U each of Exonuclease I (NEB M0293) and Exonuclease III (NEB M0206) in a 50uL reaction supplemented to 1X with CutSmart buffer. Ligates were column cleaned (Zymo-6), then linearized with 15U of RNase HII (NEB M0288) in a 30uL reaction in 1X Thermopol buffer for 90 minutes at 37C. RNase HII also cuts concatemers of AD4 adapters into ~60bp units, which can be removed in subsequent magnetic bead purification. Linearized inserts were purified using 20uL of Axygen magnetic beads (Axygen), supplemented to a final concentration of 17% PEG 8000 and 10mM MgCl2, followed by 3 washes with 70% ethanol and elution in 30uL of water.

#### Genomic coverage estimation

To determine the amount of adapter-ligated inserts that represent 1X genomic coverage, we prepared dilutions of 0.5 ng/ul, 0.25 ng/ul, 0.1 ng/ul, 0.05 ng/ul, and 0.025ng/uL of inserts. Each dilution was amplified with two adapter-specific primers, NJ-213 and NJ-214, with annealing at 61C and a 1minute extension as determined by a cycle test. We used Q5 High-Fidelity DNA Polymerase (NEB M0491). Amplicons were Axygen cleaned as described previously. We used 8ng/well of each amplified dilution and of NG16408 stock DNA for QPCR against the following single copy targets: ACTA1, ADM, ADAM12, AXL, CFB, DLX5, Kiss1, NCOA6, Notch2, RPP30, and TOP1. For each dilution sample, targets with a dCT >5 compared to stock genomic DNA were counted as absent.

The Poisson probability (***P***) of a genomic region being present in the library is given as ***P*** = 1 - (1 - *p*)^***XN***^, in which *p* = (insert size) / (genome size), *N* = the number of partitions of the genome for the given insert size and *X* = the intended genomic coverage. The proportion of targets present as identified by QPCR were compared to the value of ***P***. Based on this model, *P ~* 0.6 for a sample with ~1X genomic coverage. The 0.1ng/uL dilution tested positive for 6 of the 11 targets or a proportion of 0.545, representing between 0.5X and 1X coverage. In this manner, 0.2ng of inserts were determined to represent ~1X genomic coverage. Equimolar amounts of independently amplified replicates were mixed to obtain a pool of inserts at 5X genomic coverage.

#### Insert cloning and N25 barcoding of the GRAMc library

We cloned 30ng of 5X genomic inserts into the two-pieces of linearized GRAMc vector, SCP-GFP and the backbone cassettes, in a 1:1:1 molar ratio in a 16uL NEBuilder HiFi Assembly reaction (NEB E2621) for 20minutes at 50C. Assembled linear DNA was column purified and eluted in 20uL water. To prepare the assembled library for barcoding, 4 replicates of 8ng of the purified assembly were amplified in 9 cycles of PCR, as determined by a cycle test, with primers NJ-101 and NJ-126 using an annealing temperature of 62C and a 5 minute extension time. The replicates were combined and column-cleaned.

To add N25 barcodes downstream of the GFP ORF, we used 150ng of the library for a single cycle of PCR with NJ-127, which contains random 25bp barcode sequences, core Poly(A) signal ^4^ and 5’ biotinylation, in a 50uL Q5 High-Fidelity DNA Polymerase reaction with an annealing temperature of 60C for 40 seconds and an extension time of 15 minutes. We used NJ-126 as a competitor in the PCR to reduce the potential for template switching by occupying and extending the opposing strand. Primers were removed by Axygen bead purification using 50uL of beads and 20uL water elution, as described previously. The barcoded library was isolated using 20uL of Dynabeads MyOne C1 beads (Invitrogen 65001) with bead preparation, binding, and washing according to the manufacturer’s protocol.

Following isolation, C1 beads were washed in 20uL of water then resuspended in 50uL of water. Half of the barcoded library was amplified in 24×20uL replicate Q5 High-Fidelity DNA Polymerase reactions for 9 cycles, as determined by a cycle test, with NJ-128 and NJ-129 with 61C annealing and a 5 minute extension. Replicates were combined and Axygen bead cleaned, then gel purified (Zymo Research) with an additional Axygen bead cleaning.

Barcoded GRAMc library was then self-ligated. To reduce intermolecular ligation, we ligated 125ng of the barcoded library in 600uL of 1X T4 ligase buffer (NEB B0202) with 14,000U of high concentration T4 DNA Ligase (NEB M0202T) for 4hrs at 20C. Ligation products were supplemented with 67uL of Lambda Exonuclease buffer and 30U each of Exonuclease I (NEB M0293) and Lambda Exonuclease (NEB M0262S) for 1hour at 37C, then spiked with 1uL of ProteinaseK (ThermoFisher) for 15 minutes at 37C. Proteinase K treatment reduces viscosity of the ligation mix and increases DNA yield by nearly two fold. The library was purified with 25uL of magnetic beads (Axygen) supplemented to a final concentration of 15% PEG 8000 and 10mM MgCl2, followed by 4 washes with 70% ethanol and elution in 6.5uL of water. The product of this process is a pure population of circularized GRAMc library.

#### Transformation and size estimation of the GRAMc library

To determine the scale of electroporation, 1ul of ligation product was electroporated into 25uL of ElectroMAX DH10B competent cells (ThermoFisher 18290015). Transformants were resuspended into 1ml of pre-warmed SOC media immediately, and we used 1/500th of the transformants for 10-fold serial dilution and plating without recovery to estimate the number of colonies for the entire pool. The scale of transformation to reach the target colony number is determined based on this test. Electroporation of 4 - 10ng of ligation products generates ~40M colonies.

To generate a full GRAMc library with a colony target of 200M, duplicate electroporations were performed using 30ng of library ligates (12ng/u) per each of 2×25uL of ElectroMAX DH10B competent cells. Each replicated was resuspended into 1ml of SOC media immediately following electroporation and then replicates were combined. We used 1/2000 of the transformants for 10-fold serial dilution and plating without recovery to estimate the size of the GRAMc library. The remaining transformants were immediately used to inoculate 180ml of LB to which 100ug/ml Ampicillin was added following a 20minute recovery followed by overnight culturing. The plasmid library was prepared using the ZymoPure II Plasmid Maxiprep Kit (Zymo Research). Hereafter, this library is called Hs800_GRAMc library.

As a quality control step, twelve colonies from the plate were picked, and plasmids were extracted to check insert sizes and barcodes by Sanger sequencing. Plasmids from each colony should contain an insert (~800bp) and a barcode. Note that, due to the high barcode diversity in the ligation product, the barcode sequences identified from colonies should not be present in the final library.

The sequences of GRAMc vector and oligomers used are available in **Supplementary File 6**.

### 2. GRAMc library characterization by Illumina Paired-end sequencing

#### Sequencing library

To identify inserts and associated barcodes in individual reporter constructs, we used paired-end sequencing on the NextSeq500 platform. Two critical problems to sequence the Hs800_GRAMc library on the Illumina platform were *i*) the length of reporter constructs was too long for paired-end sequencing and *ii*) lack of diversity in the adapter sequences is incompatible with Illumina platform. To solve the length problem, we reduced the length of constructs by brining inserts and N25 barcodes closer by deleting either SCP-GFP region or the vector backbone by inverse PCR and self-ligation. To solve the low sequence diversity problem, we used a set of phased primers ^5^ to artificially increase sequence diversity. Generation of two different populations of sequencing libraries that lack either SCP-GFP region or the vector backbone also increases sequence diversity at the adapter region (see below) (**Supplementary Fig. 3**).

Sequencing library building starts with cutting 500ng of the maxi-prepped plasmids with Cas9 (NEB M0386) using sgRNAs against either the vector backbone or the GFP ORF. Both sgRNAs were predicted to have 7 off-targets in the human genome (http://crispr.mit.edu/). Primer pairs, NJ-179/NJ-183 and NJ-180/NJ-183, were used to produce templates for *in vitro* transcription of sgRNAs that respectively target backbone and GFP. The sequences of primers are available in **Supplementary File 6**. The CRISPR cut plasmid libraries were mixed with an equimolar amount of uncut plasmid libraries. Inverse PCR of 5ng of the GFP-cut linear library mixture was performed using NJ-209 and NJ-141 (denoted as “Hs800_23”) to remove the SCP-GFP region, and inverse PCR of 5ng of the backbone-cut linear library mixture was accomplished using NJ-208 and NJ-142 (denoted as “Hs800_14”) to remove the vector backbone. We used Q5 High-Fidelity DNA Polymerase (NEB) for PCR. A total of 20 replicates were prepared per template/primer pairs. Respective replicates were combined, column concentrated, gel isolated and Axygen bead cleaned. Respective amplifications were self-ligated at a concentration of 75ng in 350uL of 1X T4 DNA Ligase buffer with 3uL of concentrated T4 ligase overnight at 20C, supplemented with 20U each of Exonuclease I and Exonuclease III at 37C for 1hr, followed by incubation with Proteinase K for 10 minutes at 37C. Ligates were Axygen bead cleaned and eluted in 30uL of water.

To amplify insert::N25 cassettes, from the circularized first round PCR products, 4 replicates containing 2ng of Hs800_14 ligates were amplified using NJ-209 and NJ141 (now denoted as Hs800_1423), and 4 replicates containing 2ng Hs800_23 ligates were amplified using NJ-208 and NJ142 (now denoted as Hs800_2314) with an annealing temperature of 60C and an extension time of 90 seconds for a total of 8 cycles. Products were column cleaned, gel isolated, and bead cleaned for subsequent PCR amplification to add PE adapter sequences for Illumina sequencing.

To increase diversity of the Hs800_1423 and Hs800_2314 sequencing libraries for sequencing on the Illumina platform, each library (Hs800_1423 and Hs800_2314) was amplified using 7 different, phased PE1 containing primers. For the Hs800_1423 library, 2ng of template were used per each separate reaction with the PE2 containing primer NJ-401 and each of the following partial PE1 containing primers: NJ-400, NJ-504, NJ-505, NJ-506, NJ-507, NJ-508, and NJ-509 with an annealing temperature of 60C and an extension time of 90 seconds for a total of 7 cycles. For the Hs800_2314 library, 2ng of template were used per each separate reaction with the PE2 containing primer NJ-403 and each of the following partial PE1 containing primers: NJ-402, NJ-498, NJ-499, NJ-500, NJ-501, NJ-502, and NJ-503 with an annealing temperature of 60C and an extension time of 90 seconds for a total of 7 cycles. (We later found that the phased PE1 primers can be pooled before PCR amplification to simply the procedure.) Individual amplifications were column cleaned, gel isolated, and Axygen bead cleaned. Each of the 7 phased Hs800_1423 libraries was amplified using NJ-497 and NJ-401 to complete the PE1 adapter sequence. Each of the 7 phased Hs800_2314 libraries were amplified using NJ-497 and NJ-403 to complete the PE1 adapter sequence. For each amplification, 2ng of respective library templates were amplified in 6 cycles of PCR with an annealing temperature of 60C and an extension time of 90 seconds. Libraries were again purified, gel isolated, and Axygen bead cleaned. Equimolar amounts of the 14 phased libraries (7 from each direction) were combined to the 90% of the sequencing pool plus 10% PhiX control and used for Paired-end sequencing. The sequences of primers are available in **Supplementary File 6**.

#### Trimming adapter sequences from inserts and barcodes

The 5’- and 3’-ends of an insert and its associated N25 barcode were extracted from each pair of sequence reads. We used Trimmomatic ^6^ to remove adapter sequences and seqtk (https://github.com/lh3/seqtk) to reverse complement sequences. To extract the 5’-end and 3’-end of an insert, we trimmed P1 and P2 adapters, respectively. To extract N25 barcodes, depending on the orientation of a sequence read, we trimmed P3 or P4 adapter first, reverse complemented the trimmed sequence, and trimmed P4 or P3 adapter. Paired-end reads that failed to trim any adapter sequence were abandoned. Note that in the case of N25 barcode sequences, we kept 1bp from each adapter, resulting in 27bp reads. Adapter sequences used for trimming are available in **Supplementary File 6.**

#### Mapping sequence reads and identification of inserts in the human genome

To identify inserts, extracted 5’- and 3’-ends of inserts were mapped on to the GRCh38/hg38 assembly (downloaded from genome.ucsc.edu). We used Burrows-Wheeler Alignment tool (BWA) ^7^ to map sequences with the following command: “bwa mem -W 1500.” Mapped pairs of reads that spanned >1,500bp or <300bp were abandoned. When two mapped inserts overlapped and their mid-points were within 20bp range and both ends within 50bp range, we combined them into one insert taking the coordinates that maximize its length.

#### Clustering N25 barcodes

To identify reads from the same barcode, we clustered the extracted barcode reads based on the following procedure: *i*) Representative reads were generated by filtering redundant reads by using the Khmer software package ^8^ with the command: “normalize-by-median.py -C 1 -k 25 -N 5 -x 2.5e9.” *ii*) The entire set of barcode reads were matched against the representative reads using the BWA software ^7^ with the command: “bwa aln -n 2 -o 2 -e -1 -M 3 -O 11 -E 8 -k 1 -l 6.“ Barcode reads that did not match any of the representative reads were added to the representative reads file and repeated the BWA search. Reads for the same barcodes were identified by single-linkage-clustering, and each cluster was assigned a unique barcode cluster (bcl) number. A new file of representative reads with the bcl numbers was generated for future use (See 3. GRAMc assay in HepG2: *Matching barcode reads to barcode clusters*).

#### Associating genomic inserts with barcode clusters (bcls)

Although each barcode read is inherently connected to reads from an insert in paired-end reads, we have observed that a minor fraction of bcls were associated with more than one genomic inserts identified in the above. The main reason for this ambiguity was due to highly similar duplicated regions in the genome. We forced the assignment of a bcl to an insert that has most reads for the bcl. In case ≥2 inserts had the same number of reads for a bcl, we did not assign the bcl to any insert.

### 3. GRAMc assay in HepG2

#### Cell culture

HepG2 cells (ATCC HB-8065) were grown under supplier recommended conditions of EMEM supplemented with 10% fetal bovine serum without antibiotics. HepG2 cells were used within no more than 16 passages from receipt for all experiments. All experiments were performed in cells that underwent a minimum of 5 passages from thawing, because reporter expressions in cells of <5 passages vs. cells of ≥5 passages were very different.

#### Genome-scale transfection and lysate collection

For each genome-scale transfection batch, 10^7^ cells were seeded in 30ml media in each of 10×150mm culture dish (100M cells) and allowed to attach for 30 hours. Cells were transfected with 100ug of the Hs800_GRAMc library using 100uL of DNA-IN for HepG2 reagent (MTI-Globalstem) in 4ml of Optimem (ThermoFisher) prepared in 2×2-ml siliconized tubes according to the manufacturer’s protocol. A total of 10 10×150mm dishes were used to collect ~200M cells per batch.

For collection, cells were washed with 1X PBS 26hours after transfection and were collected by scraping in 2.4ml RNA-STAT-60 (AMSBIO) per plate. Lysates were combined and prepared according to the manufacturer’s protocol with the addition of a second 70% ethanol wash.

#### RNA preparation and cDNA synthesis

Our protocol focuses on two key parameters: *i*) comprehensively removing contaminated DNAs in RNA sample and *ii*) maximizing the efficiency of reverse transcription (RT) with a large quantity (~4mg) of total RNA. We have achieved the first goal by complementing DNase I with a cocktail of Exonuclease I and III to comprehensively remove both double stranded and single stranded DNAs, because DNase I is less efficient against single stranded DNA. To achieve the second goal in a cost-efficient way, we use 15 times more RNA than manufacturer’s recommended maximal input RNAs without compromising cDNA yield in RT reaction. A schematic of the procedure is available in **Supplementary Figure 4**.

To remove contaminated DNAs, isolated total RNA (~4mg) was resuspended in 1.7ml of nuclease free water digested for a minimum of 4 hours at 37C in a 2ml reaction containing 1X DNase I Buffer, 100U of DNase I (NEB M0303), and 900U each of ExoI and ExoIII. The progress of DNA removal was monitored by QPCR against the GFP ORF (NJ-443 and NJ-444). For this quality control step, a diluted sample of RNA was heat inactivated at 80C for 20 minutes and loaded at an equivalent volume of ~1000cell/well. As needed, DNase digestion was allowed to proceed overnight until QPCR C*t* value becomes greater than 30. Following digestion, nucleases were removed by extraction with Phenol:Chloroform:Isoamyl alcohol (25:24:1) and ethanol precipitated overnight at −20C followed by two washes with 75% ethanol. RNA was resuspended in 1ml of RNase-free water.

As a quality control for reverse transcription (RT), an equivalent volume of the total RNA containing ~4000cells (~1ug) was used for cDNA synthesis using the High Capacity cDNA Reverse Transcription Kit (Applied Biosystems 4368813) following the manufacturers protocol with the addition of 5pmole of a GRAMc library specific RT oligo (NJ-489) and used as the standard for maximum cDNA synthesis from transcripts.

The remaining total RNA (~4mg) was diluted to 1.420ml, and 2000pmol of GRAMc_RT_oligo (NJ-489) was added. The RNA/primer mixture was incubated at 65C for 1 minute and chilled on ice, followed by addition of 200uL of 10x High Capacity buffer, 80uL of 10mM dNTP and 100uL of Multiscribe without using random oligomers. The reaction was incubated for 10 minutes at room temperature, and then 4 hours at 37C. We monitored progression of the genome-scale cDNA synthesis via QPCR against GFP in comparison to the standard RT control using an equivalent volume of 100cells/well. Reactions were allowed to proceed until C*t* value becomes similar to that of the standard RT reaction. If needed, reactions were spiked with M-MuLV Reverse Transcriptase (NEB M0253) and additional dNTPs and allowed to proceed overnight.

Upon completion of the RT reaction, the samples were ethanol precipitated to reduce the volume. RNA/cDNA was resuspended and digested with 1000U of RNase If (NEB M0243) in a 500uL reaction with 1X NEB3 buffer at 37C overnight. For removal of excess protein, 1uL of Proteinase K solution was added to the reaction and incubated at 37C for 15 minutes. cDNA was ethanol precipitated overnight at −20C with glycogen as carrier, washed 3x with 80% ethanol. cDNA pellets were resuspended in 200uL of water and heated to 95C for 10 minutes to destroy residual Proteinase K. A sample of the cDNA library was subjected to quality control by QPCR.

#### Preparation of expressed N25 barcodes for NGS

The entire pool of expressed N25s were amplified using primers NJ-141 and NJ-142 in 8 replicates of a 50ul Q5 PCR reaction using an annealing temperature of 62C and an extension time of 1minute for a total of 8 cycles. Replicates were combined for each batch. A 50uL aliquot was processed from each batch as follows: unwanted long DNAs were bound using a 0.5X volume of Axygen beads for 20 minutes at room temperature. The desired short amplicons (65bp) from the supernatant were further purified for each batch using duplicate Zymo column and eluted each in 20uL of water. To prepare amplicons for sequencing expressed barcodes, 2ng of 1^st^ round amplified and cleaned N25 barcodes were subjected to another 9 cycles of amplification with NJ-141 and NJ-142. To prepare amplicons for sequencing the input library, 2ng of the input library were amplified in 9 cycles of PCR from a mixture of uncut/CRISPR backbone cut/CRISPR GFP cut plasmid library template using the NJ-141 and NJ-142 primers.

Sequencing libraries were prepared both for IonTorrent Proton sequencing (Batch 1: NJ197 and NJ-523; Batch2: NJ-198 and NJ-523) and Illumina NextSeq500 sequencing (14 phased libraries using NJ-400/NJ-504/NJ-505/NJ-506/NJ-507/NJ-508/NJ-509 with NJ364 or NJ-402/NJ-498/NJ-499/NJ-500/NJ-501/NJ-502/NJ-503 with NJ-399) ^5^. For all of these amplifications, an annealing temperature of 65C and an extension time of 20seconds were used for a total of 6 cycles.

The sequences of primers are available in **Supplementary File 6**.

#### Matching barcode reads to barcode clusters (bcls)

The goal of this step is to count the number of barcode reads from either expressed barcodes or the input library for each barcode cluster (bcl). Adapter trimmed barcode reads were matched to the representative barcode reads established in the above by using BWA search with the same command as above. When a barcode read matched more than one bcl, we counted each match to the respective bcls. Because the same procedure was applied to both expressed barcodes and the input library, the effect of multiple counting of a barcode read is neutralized.

#### Computation of CRM activity

The goal of this step is to compute *cis*-regulatory activity of each insert based on the number of reads for each bcl that are counted from expressed barcodes and the input library. When an insert is associated with ≥2 bcls (99% of inserts), we combined the read counts for all bcls for the insert. First, to avoid falsely calling CRMs due to too low input counts, we retained inserts with ≥10 counts from the input library or ≥50 counts of expressed barcodes for both batches of experiments. This filtering resulted in 9,339,996 inserts that meet the retention criteria. Second, read counts for expressed barcodes were divided by the read counts for the input library, and the resulting numbers were rank ordered. We used the middle 30% of data to compute the background activity (bg) following our earlier approach in sea urchin embryo ^9^. CRM activities are further normalized to the background activity. An insert was called CRM, when at least one batch showed ≥5xbg and another ≥4.5xbg (90% of 5xbg). We have identified 54,115 inserts that passed the criteria. After removing inserts that have ≥95% identical sequences in other part of the genome and merging overlapping CRMs, the final set contains 41,216 unique and non-overlapping CRMs.

A scatter plot shown in **Figure 2a** was generated by using ggplot2 ^10^ in the R package (cran.r-project.org) using 500,000 randomly selected inserts.

### 4. Genomic distribution of CRMs

To compare genomic locations of CRMs and genes, we used publicly available gene annotation file “GRCh38.89.gff3” from ftp.ensembl.org and RNA-seq data for HepG2 cells “ENCFF861GCR and ENCFF640ZBJ” from encodeproject.org. Genes with FPKM ≥1 in both RNA-seq data were considered “expressed” in this paper. To generate maps shown in **Figure 2c** and Supplementary **Figure 5**, we used the Grid Graphics Package ^11^ in R with the bin size of 1Mb.

To compute enrichment of CRMs in genomic regions with respect to genes (**Fig. 2d**), insert/CRM that span more than 2kb window were assigned to a window that overlaps most with the insert. Genomic coordinates of the 5’-end and 3’-end of a gene were extracted from GRCh38.89.gff3 file. An insert/CRM was counted only once for a gene, but was allowed to be counted multiple times for different genes.

### 5. One-by-one reporter assay for validation

#### Making individual reporter constructs

The 21 genomic regions (11 CRMs, 5 marginally active regions and 5 inactive regions) were individually amplified by PCR and cloned into a pre-barcoded SCP-GRAMc vector ^12^ by Gibson Assembly ^13^. Primers used to amplify inserts contain flanking sequences that overlap with adapter sequences present in the vector. Each assembly was done in a 2uL NEBuilder HiFi Assembly reaction. Assembly reactions were used to transform Mix and Go DH10B competent cells (Zymo Research T3019), and positive clones were identified by colony PCR. Endotoxin-free plasmids were prepared (Zymo Research D4208T).

The pre-barcoded SCP-GRAMc vector was further used to generate an EGFP internal control vector for use in QPCR of GFP reporter expressions for individual clones. For this the vector was amplified by inverse PCR with NJ731 and NJ732. The EGFP ORF from pEGFP-C1 was amplified using NJ729 and NJ730 and Gibson assembled to the SCP-GRAMc vector at a ratio of 2:1 using NEBuilder HiFi Assembly master mix. Note that the GFP ORF used in the GRAMc vector is different from the commonly used EGFP ORF, and the two GFPs can be differentially detected by QPCR.

The sequences of primers are available in **Supplementary File 6**.

#### Individual reporter assay to validate GRAMc results

HepG2 cells were seeded at ~60K cells per well of a 24-well plate in 500uL of EMEM supplemented with 10% FBS. For consistency with the genome-scale assay, cells were used between passages 12 and 15 from receipt from ATCC and at least 7 passages after recovery. Cells were allowed to attach for 24-hours and transfected with a mixture of 50uL Optimem, 200ng of GFP containing individual test plasmids, 200ng of SCP-EGFP control vector, and 1.2uL DNA-In reagent. After 26 hours (~80-85% confluency, in keeping with the genome-scale assay), cells were washed twice in DPBS, collected in 300uL of DNA/RNA lysis buffer (ZymoResearch), and gDNA and total RNA for each sample were purified using Zymo II columns with binding and washing as per the manufacturer’s protocol. RNA was eluted in 34uL of water. Half of the total RNA for each sample was treated in a 20uL Turbo DNase reaction (ThermoFisher) for 1hour at 37C. The reactions were terminated with 2uL of DNase inactivation reagent (ThermoFisher). Half of the DNase treated RNA was used in a 20uL 1X High-Capacity cDNA synthesis reaction with an additional 10pmole of GRAMc_RT_oligo (NJ-489) and RNase inhibitor. QPCR was performed against GFP and EGFP on a total gDNA equivalent of 1/40,000 of the original sample, a non-RT control equivalent of 1/40 of the total RNA sample, and a cDNA equivalent of 1/160 of the original sample. GFP expressions driven by individual test fragments were normalized to the internal control (EGFP expression, NJ404/NJ405). The sequences of QPCR primers are available in **Supplementary File 6**.

### 6. Relative enrichment of ENCODE annotations in CRMs vs. inactive inserts

ENCODE ChIP-seq files were obtained from encodeproject.org. The list of files is available in **Supplementary Table 3**. Overlap between CRMs and individual ENCODE data was computed using bedtools ^14^ with the command: “bedtools jaccard -f 1E-09 -F 1E-09.” The relative enrichment of ENCODE annotations in CRMs was computed in the following procedure: *i*) We first computed the genomic proportion of overlapping base pairs between CRMs and an ENCODE annotation. *ii*) We computed randomly expected overlap by multiplying genomic proportions of the two datasets. *iii*) Result from *i*) is divided by result from *ii*) to compute the enrichment. *iv*) Following the same procedure, we computed enrichment of the same ENCODE annotation in inactive regions (L1 group). *v*) Relative enrichment was computed by taking the ratio of *iii* and *iv*.

### 7. Motif enrichments in CRMs and predicted strong enhancers

#### Selection of GRAMc inserts

Predicted strong enhancers for HepG2 by ChromHMM ^15,16^ were compared to GRAMc data in terms of CRM activity and motif enrichment. Genomic coordinates of chromatin states were converted via liftOver ^17^ to hg38. We first randomly selected nonoverlapping GRAMc inserts that ≥90% overlap in its length with predicted strong enhancers. This selection process collected 18,898 GRAMc inserts that correspond to predicted strong enhancers. This data was utilized to generate **Figure 3a**. The list of selected inserts is provided as **Supplementary File 3**.

To compare motif enrichment, we randomly sampled another 18,898 nonoverlapping GRAMc CRMs (≥5xbg or G5) without considering predicted enhancers. We also sampled 37,796 nonoverlapping inactive (≤1xbg or L1) inserts as negative control.

#### Motif enrichment survey

To survey putative transcription factor binding site (TFBS) motifs, the 75,592 inserts sampled were analyzed simultaneously. We used the HOCOMOCOv10 database ^18^ and the FIMO software ^19,20^ with an E-value cutoff of 1E-5. Abundance of each motif is the proportion of motif-harboring inserts for a given set. Relative motif enrichment was computed by dividing the abundance of a motif in CRMs or predicted enhancers by the abundance of the same motif in the negative control set (inactive inserts).

#### Comparison of enrichments of motifs and ChIP-seq peaks in CRMs

Fifty-eight common transcription factors between the HOCOMOCOv10 and the ENCODE ChIP-seq data were identified by their names. Relative enrichment scores computed in the above were used to generate **Figure 4b**.

### 8. Measuring the effect of gene ectopic expression on CRMs

#### Preparation of random sub-sets of the GRAMc library

To obtain small-scale subsets of the GRAMc library for perturbation experiment by ectopic expression of *pitx2* or *ikzf1*, ~50uL of frozen glycerol stock was diluted into 2ml of LB media, recovered with orbital shaking 250 RPM at 37C for 20 minutes. We prepared a series of 2-fold dilutions, 1/100^th^ of which was used for 2 10-fold dilutions for plating and colony counting, and the remainder of each 2-fold diluted culture was used to seed 150ml LB-Amp cultures for overnight growth as described previously. Cultures that were estimated to contain ~80,000 colonies (80K library) were processed using the ZymoPure Plasmid Maxiprep Kit.

#### Perturbation assay of the 80K construct library

Cells were seeded in duplicates of ~2M cells per 10 cm^2^ plate for transfection with each of 3 cotransfections: 80K library + CMV::pitx2 (Genscript OHu17480D), 80K library + CMV::IKZF1 (Genscript OHu28016D), and 80K library + CMV::EGFP (Clontech pEGFP-C1). Cells were cultured for ~24h prior to transfection. Cells were co-transfected with 9ug of the 80K library and 3ug of the respective expression vector using 36uL of DNA-IN for HepG2 reagent (MTI-Globalstem) and 1.2ml of Optimem (ThermoFisher) prepared according to the manufacturer’s protocol.

Cells were harvested by trypsinization and washing with 1X DPBS 24h after transfection. A 1/10^th^ portion of the cells was saved for Western Blot analysis to confirm expression of Pitx2 and IKZF1. The remaining cells were lysed and processed using the Zymo-Duet kit with the IIICG column for both DNA and RNA, without on-column DNase I treatment. DNA was eluted in 100uL and RNA was eluted in 80uL and treated with DNase I (8U)/ExoI(100U)/ExoIII(100U) for a minimum of 4hr at 37C in a total reaction volume of 100uL in 1X DNase I buffer. Assuming ~10M cells per sample, an ~10,000 cell equivalent of gDNA and an ~5000 cell equivalent of nuclease treated RNA were tested with QPCR with GFP as target to confirm the quality of transfection and completion of DNA removal in RNA, respectively. Reactions were spiked with another 2U of DNase I as needed. RNA was column cleaned using a Zymo-IIIC column and eluted in 50uL of water. An equivalent of ~4000cells was used as a measure of quality control in a standard RT reaction as described in the genome-scale protocol. The remaining RNA was incubated with 80pmole of GRAMc_RT_oligo (NJ-489) used for cDNA synthesis in an 80uL 1X High-Capacity cDNA synthesis reaction using 8uL of Multiscribe and 3.2uL of dNTP, but without the use of random primers, for 4hrs to overnight at 37C as per a quality control QPCR following 2hrs of RT. Upon completion of DNA digestion, 4uL of NEB3 buffer and 2uL of RNase If were added to the reaction for 2hr at 37C, then spiked with Proteinase K for 15min at 37C, and heat inactivated for 10min at 95C followed by overnight ethanol precipitation and resuspension in 30uL of water.

N25 barcodes were preliminarily amplified as described previously, but using 6cycles of a single 50uL Q5 High-Fidelity DNA Polymerase reaction, and IX barcoded for IonTorrent Proton sequencing using the following primer pairs: For control-1: NJ-197/NJ523, for control-2: NJ-198/NJ523, for Pitx2-1: NJ-200/NJ523, For Pitx2-2: NJ-132/NJ523, for IKZF1-1: NJ-133/NJ523, for IKZF1-2: NJ-134/NJ523.

Data analysis was conducted as described above. The sequences of primers are available in **Supplementary File 6**.

#### Confirmation of ectopic Transcription Factor expression by Western Blot

An aliquot of each transfection condition (80K library + CMV::pitx2, 80K library + CMV::IKZF1, and 80K library + CMV::EGFP) was lysed in 80uL of RIPA buffer (150mM NaCl, 1% NP40, 0.5% Sodium Deoxycholate, 0.1% SDS, 50mM Tris-HCl pH 8.0, 5mM EDTA) spiked with a 1:100 dilution of Halt Protease Inhibitor Cocktail (Thermofisher) on ice for 30min with intermittent flicking. Lysates were centrifuged at 12,000RPM for 10 min at 4C and quantified using BCA reagent.

Approximately 25ng of each sample was loaded in duplicate sets (expressed and control), separated on a 12% polyacrylamide gel, transferred to a PVDF membrane, and blotted with antibodies against FLAG (1:500, Santa Cruz sc-166355) or GAPDH (1:1000, Santa Cruz sc-25778). Horseradish peroxidase-conjugated secondary antibodies (1:5000) and enhanced chemiluminescence reagents (GE Healthcare) were used to detect bands on a Bio-Rad ChemiDoc MP system.

### 9. Enrichment of repeat elements in CRMs

The list of annotated repeat elements was downloaded from http://www.repeatmasker.org/genomes/hg38/RepeatMasker-rm405-db20140131/hg38.fa.out.gz ^21^. The following procedure describes how the fold-enrichments of repeat elements in different cohorts of GRAMc activity were computed. First, the overlap between the GRAMc-tested inserts and individual repeat families were detected using bedtools ^14^ with the command: “bedtools intersect -wao -a”. Second, the proportion of each repeat family in each cohort of GRAMc-activity was computed by dividing the cohort-specific number of inserts that overlapped with the repeat family by the total number of inserts that overlapped with the same repeat family. Third, the expected proportion of each cohort is computed by dividing the number of inserts in each cohort by the total number of inserts (without considering overlap with repeat elements). Fourth, the fold-enrichment of a repeat family in each cohort is computed by comparing results from steps 2 and 3. Statistical significance of enrichment or depletion of a repeat family in the G5 set was computed by the two-tailed binomial test after correcting for multiple tests.

## Supplementary Table

**Supplementary Table 1.** Reported accuracy of CRM predictions.

| Selection criteria | Number of regions considered | CRMs/regions tested | Test bed | Reference |
| --- | --- | --- | --- | --- |
| Tissue specific<br>ChIP-seq and<br>conservation | 4656 | 105/329 (32%) | Transgenic mice | 32 |
| Tissue specific<br>ChIP-seq | 2543 (Forebrain)<br>561 (Midbrain)<br>2105 (Limb) | 49/63 (78%) Forebrain<br>28/34 (82%) Midbrain<br>20/25 (80%) Limb | Transgenic mice | 33 |
| Tissue specific<br>ChIP-seq | 3597 (Heart) | 97/130 (75%) Heart | Transgenic mice | 34 |
| GWAS and eQTL<br>variants | 3642 | 2432/29173 variants (12%) | Cell lines | 35 |
| TAL1 binding | 4915 | 39/70 (56%);<br>43/66 (65%) | Cell lines;<br>Transgenic mice | 36 |
| ATAC-seq and<br>DNase-seq | 1582 (ATAC-seq;<br>PMC* to non-PMC)<br>1659 (DNase-seq;<br>PMC/non-PMC) | 9/31 (29%) | Sea Urchin<br>embryos | 37 |
| ATAC-seq and<br>FACS (endothelial<br>cells) | 5291 (Endothelial<br>cells) | 8/12 (67%) | Zebrafish<br>embryos | 38 |

\*PMC = primary mesenchyme cells

**Supplementary Table 2.** Genomic coverage of the human GRAMc library.

| **Chromosome** | **Coverage**<br>**(all)** | **Coverage**<br>**(unique, <95% identity)** |
| --- | --- | --- |
| chr1 | 0.893 | 0.860 |
| chr2 | 0.962 | 0.948 |
| chr3 | 0.970 | 0.963 |
| chr4 | 0.968 | 0.960 |
| chr5 | 0.969 | 0.940 |
| chr6 | 0.963 | 0.954 |
| chr7 | 0.963 | 0.937 |
| chr8 | 0.966 | 0.951 |
| chr9 | 0.850 | 0.804 |
| chr10 | 0.963 | 0.948 |
| chr11 | 0.961 | 0.949 |
| chr12 | 0.964 | 0.959 |
| chr13 | 0.971 | 0.951 |
| chr14 | 0.968 | 0.939 |
| chr15 | 0.970 | 0.927 |
| chr16 | 0.867 | 0.820 |
| chr17 | 0.941 | 0.902 |
| chr18 | 0.969 | 0.941 |
| chr19 | 0.921 | 0.867 |
| chr20 | 0.956 | 0.936 |
| chr21 | 0.934 | 0.827 |
| chr22 | 0.937 | 0.861 |
| chrX | 0.866 | 0.828 |
| chrY | 0.423 | 0.300 |
| **Genome** | **0.934** | **0.907** |

**Supplementary Table 3.** List of inserts for individual reporter assay.

| Cohort | Genomic Coordinate (hg38) | GRAMc activity |
| --- | --- | --- |
| G5 | chr6:1137373-1138163 | 63.43 |
| G5 | chr7:7620832-7621599 | 19.36 |
| G5 | chr1:37890375-37891109 | 13 |
| G5 | chr14:20288372-20289134 | 27.34 |
| G5 | chr15:74439985-74440809 | 211.97 |
| G5 | chr1:4465004-4465791 | 393.05 |
| G5 | chr11:41152726-41153475 | 8.68 |
| G5 | chr6:22827845-22828628 | 7.48 |
| G5 | chr8:630933-631871 | 15.73 |
| G5 | chr9:78563843-78564651 | 9.28 |
| G5 | chr12:123700613-123701394 | 6.42 |
| G3L5 | chr11:84433280-84434177 | 2.99 |
| G3L5 | chr12:2927744-2928540 | 3.65 |
| G3L5 | chr9:74920250-74921035 | 4.85 |
| G3L5 | chr5:789596-790308 | 3.2 |
| G3L5 | chr19:271467-272331 | 4.54 |
| L1 | chr1:1154867-1155707 | 0.93 |
| L1 | chr8:342822-343535 | 0.65 |
| L1 | chr10:1331187-1332088 | 0.67 |
| L1 | chr16:89802861-89803690 | 0.28 |
| L1 | chr3::45837895-45838760 | 0.29 |

**Supplementary Figure 1.**
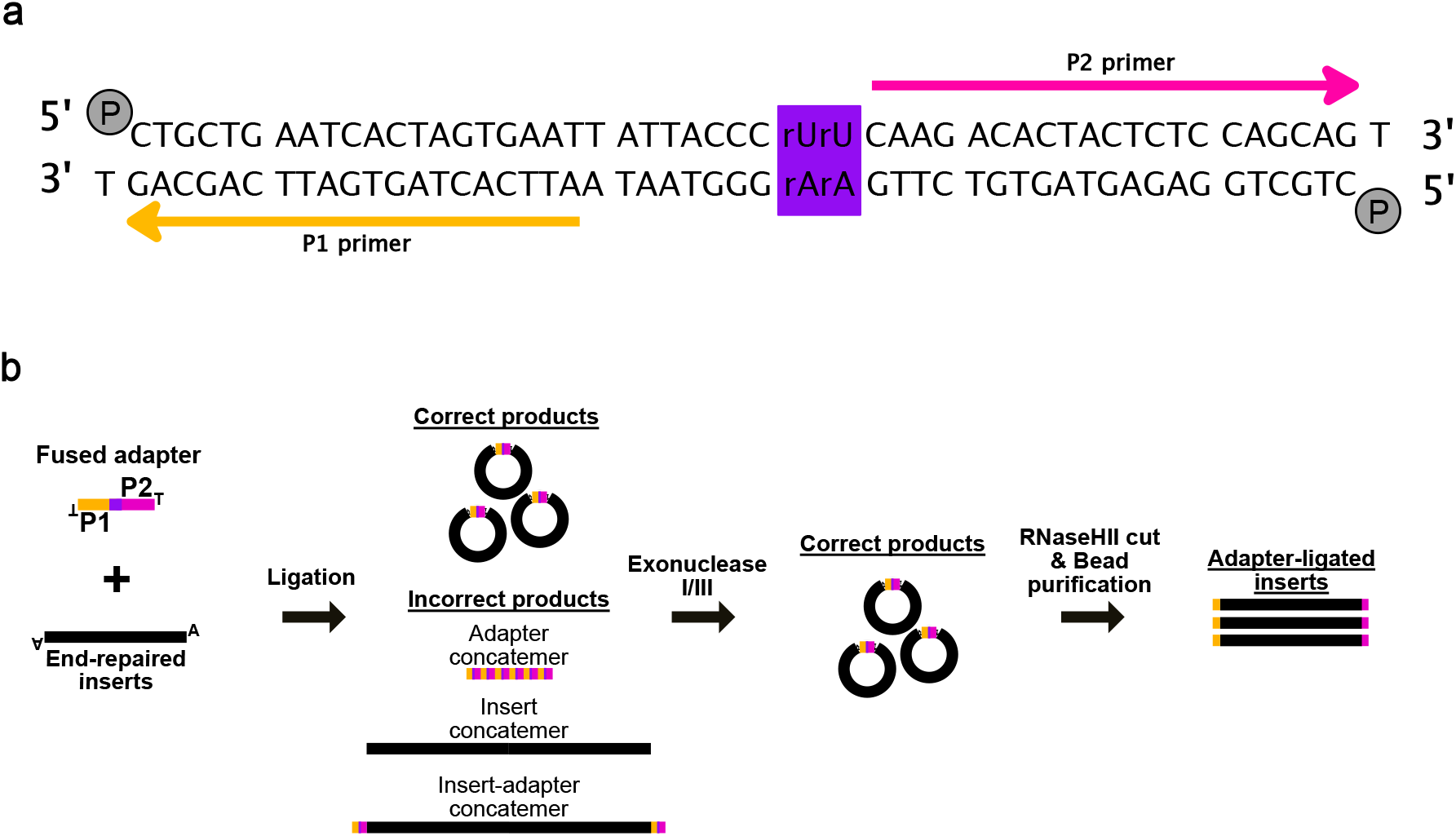
Preparation of adapter-ligated inserts with fused adapter. **(a)** Structure of the fused adapter. The fused adapter is prepared by annealing two 5’-phosphorylated oligomers. The fused adapter contains two primer sites, P1 (yellow arrow) and P2 (magenta arrow), for amplification of adapter ligated genomic inserts. The purple box indicates two ribonucleotides for RNaseHII cut. **(b)** Preparation of a pure population of adapter-ligated inserts. Ligation of an insert and a fused adapter generates a circular DNA that is resistant to exonuclease treatment. All undesired linear DNAs will be removed by exonuclease I/III. Because circular DNAs are difficult to amplify by PCR, circular ligation products are linearized by RNaseHII. Target ribonucleotide sites for RNaseHII are marked with a purple box. Linearized adapter-ligated inserts are ready for PCR amplification with P1 and P2 primers.

**Supplementary Figure 2.**
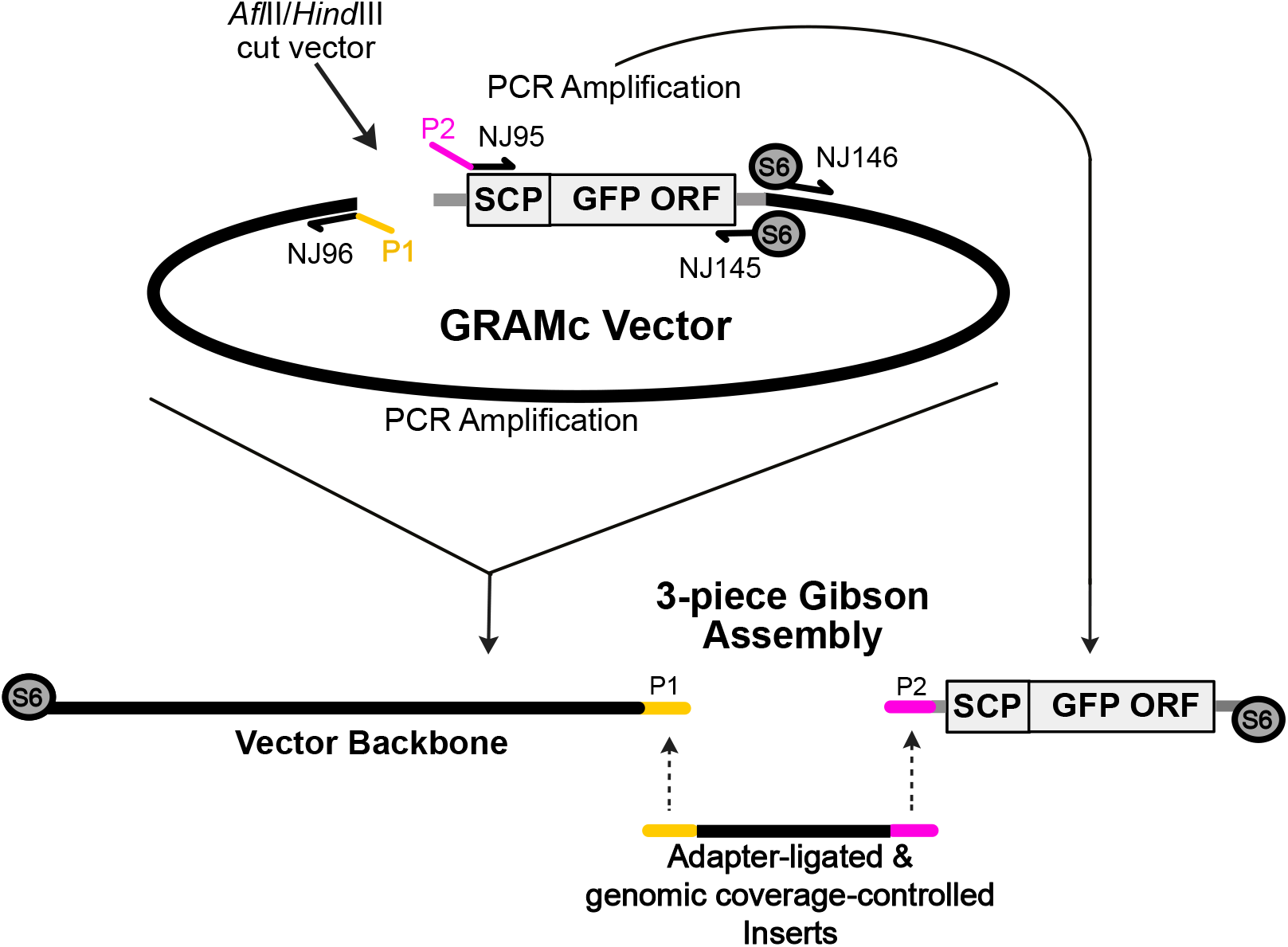
Preparation of GRAMc vector for Gibson Assembly. GRAMc vector for Gibson Assembly is prepared by PCR using two pairs of primers and *Afl*II/*Hind*III double-digested vector as template. The vector is amplified into two pieces, one containing SCP-GFP cassette with primers NJ95 and NJ145 and one containing the vector backbone with primers NJ96 and NJ146. Primers NJ96 and NJ95 respectively add the P1 and P2 adapter sites to the vector backbone and the SCP-GFP cassette, and the adapter sites are required for subsequent Gibson Assembly with adapter-ligated inserts. Primers NJ146 and NJ145 contain a sequence of 6 phosporothioated nucleotides at the 5’ end (indicated by S6) to protect the 5’-termini of PCR products from degradation during Gibson Assembly.

**Supplementary Figure 3.**
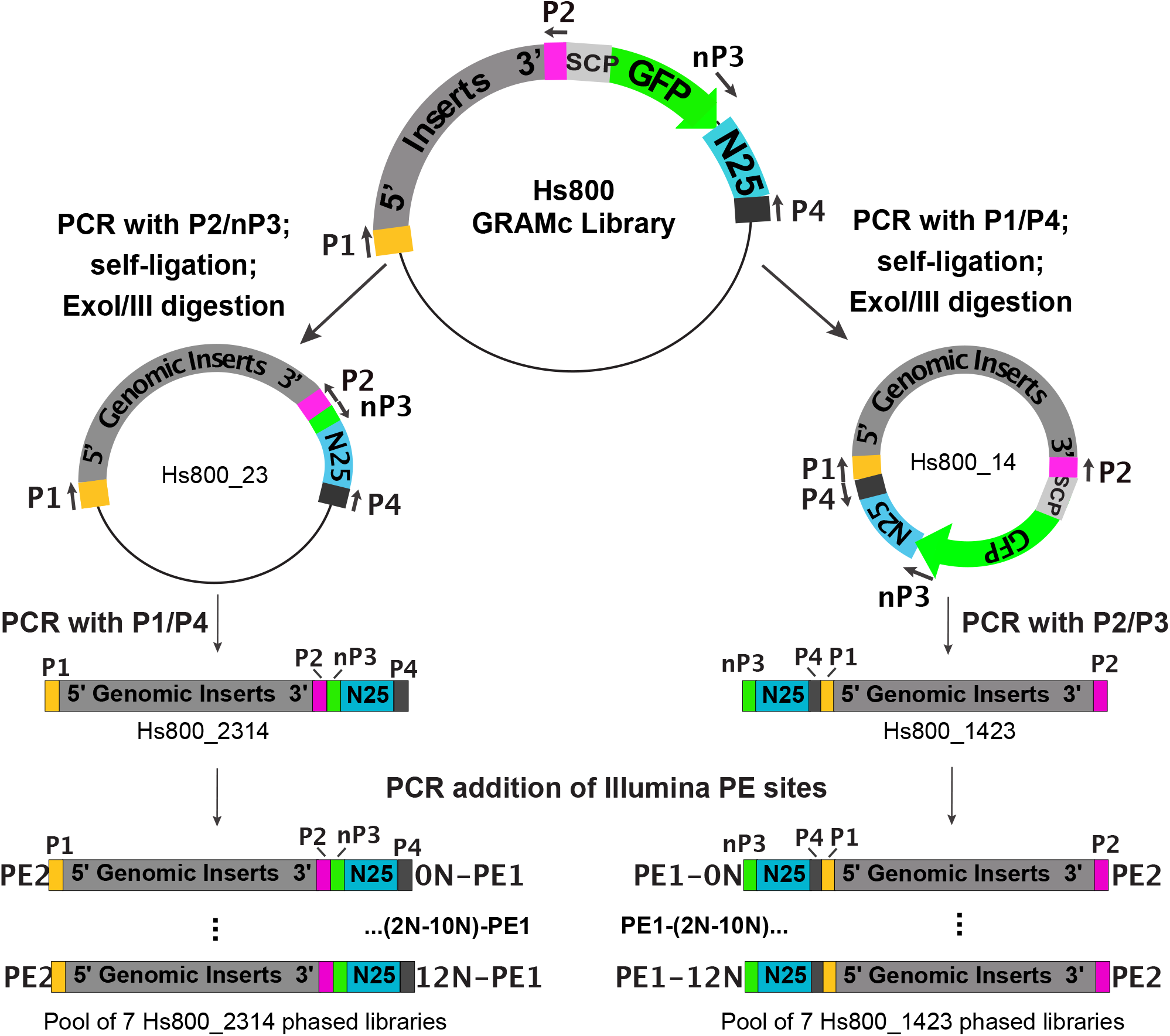
Building paired-end sequencing libraries for Illumina NextSeq500. To bring an insert and an N25 barcode in each reporter construct close to each other, the SCP-GFP cassette or the vector backbone is deleted by PCR with P2/nP3 or P1/P4 primers, respectively. Subsequent self-ligations of PCR products generate 2 sublibraries with N25s mated to either the 5’-end of inserts (Hs800_14) or the 3’-end of inserts (Hs800_23). Exonuclease treatment ensures survival of only mated circular ligates during subsequent second round amplification of insert::N25 cassettes with the alternate set of primers (P1/P4 for Hs800_23 and P2/nP3 for Hs800_14) to generate 2 sequencing libraries, Hs800_2314 and Hs800_1423. PCR adds PE1 and PE2 sites for Illumina paired-end sequencing. Addition of PE1 sites are accomplished using 7-phased primers per sequencing library to offset the lack of diversity in flanking adapter sequences. Phased primers incorporate 0N, 2N, 4N, 6N, 8N, 10N, and 12N random sequences between PE1 sites and respective nP3 or P4 sites. The 14 phased libraries are sequenced on the Illumina NextSeq500 platform.

**Supplementary Figure 4.**
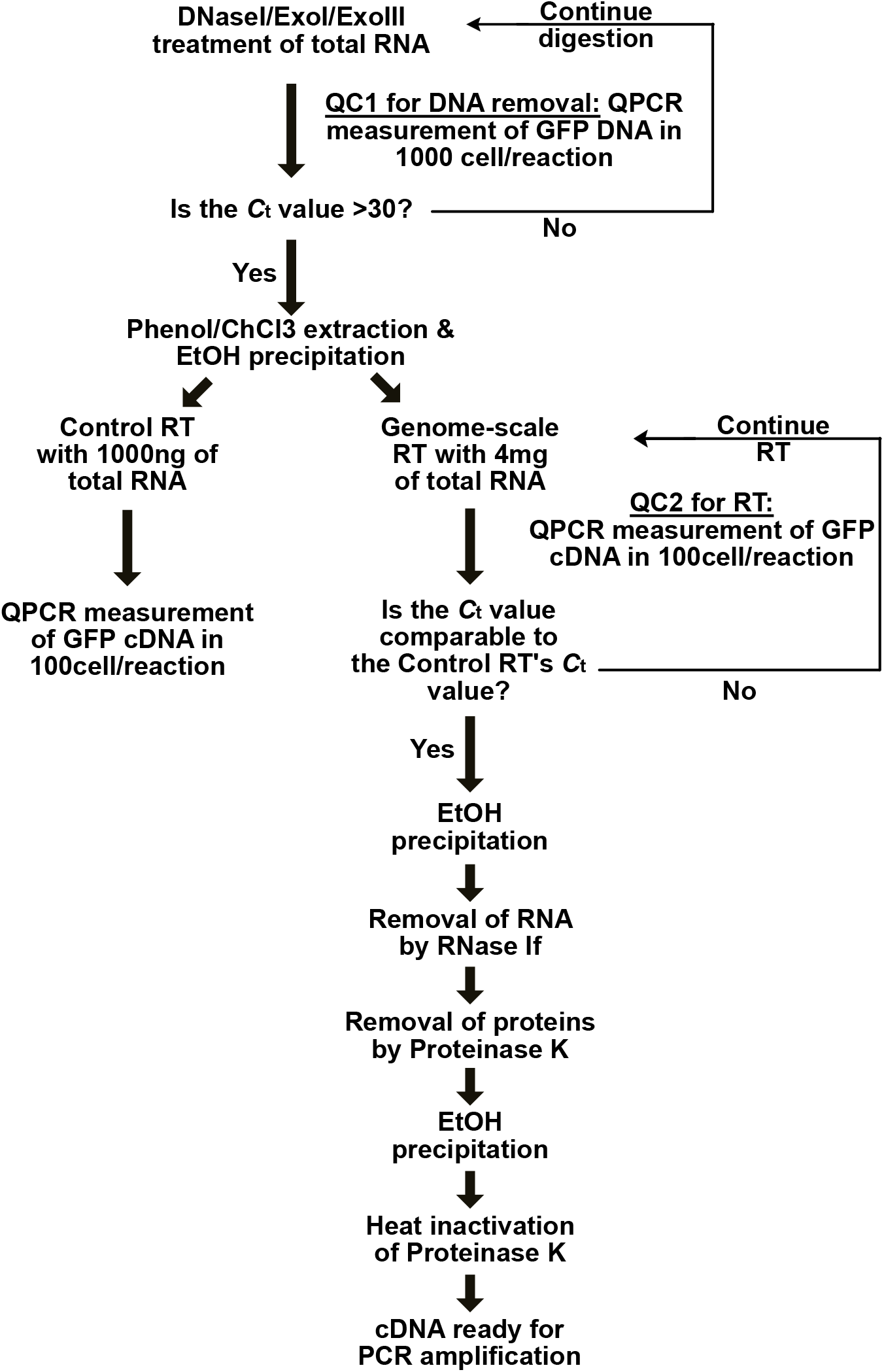
QPCR-based Quality Control (QC) steps during processing of a large quantity of total RNA into cDNA. During the first QC step (QC1), removal of contaminated DNAs in RNA samples is monitored by measuring GFP DNAs by QPCR. After 12 hours of DNase treatment, if the Ct value for GFP DNA is still ≤30, continue DNA digestion. Check the Ct value every 6 hours and repeat this process until the Ct value is >30. As a quality control (QC) standard for reverse transcription (RT), we use 1000ng of DNaseI/ExoI/ExoIII digested total RNA for standard RT reaction. During the second QC (QC2) step, the genome-scale RT reaction is monitored and supplemented with reagents as needed until the Ct value of GFP cDNA is within 1 cycle of the Ct value in the QC standard.

**Supplementary Figure 5a.**
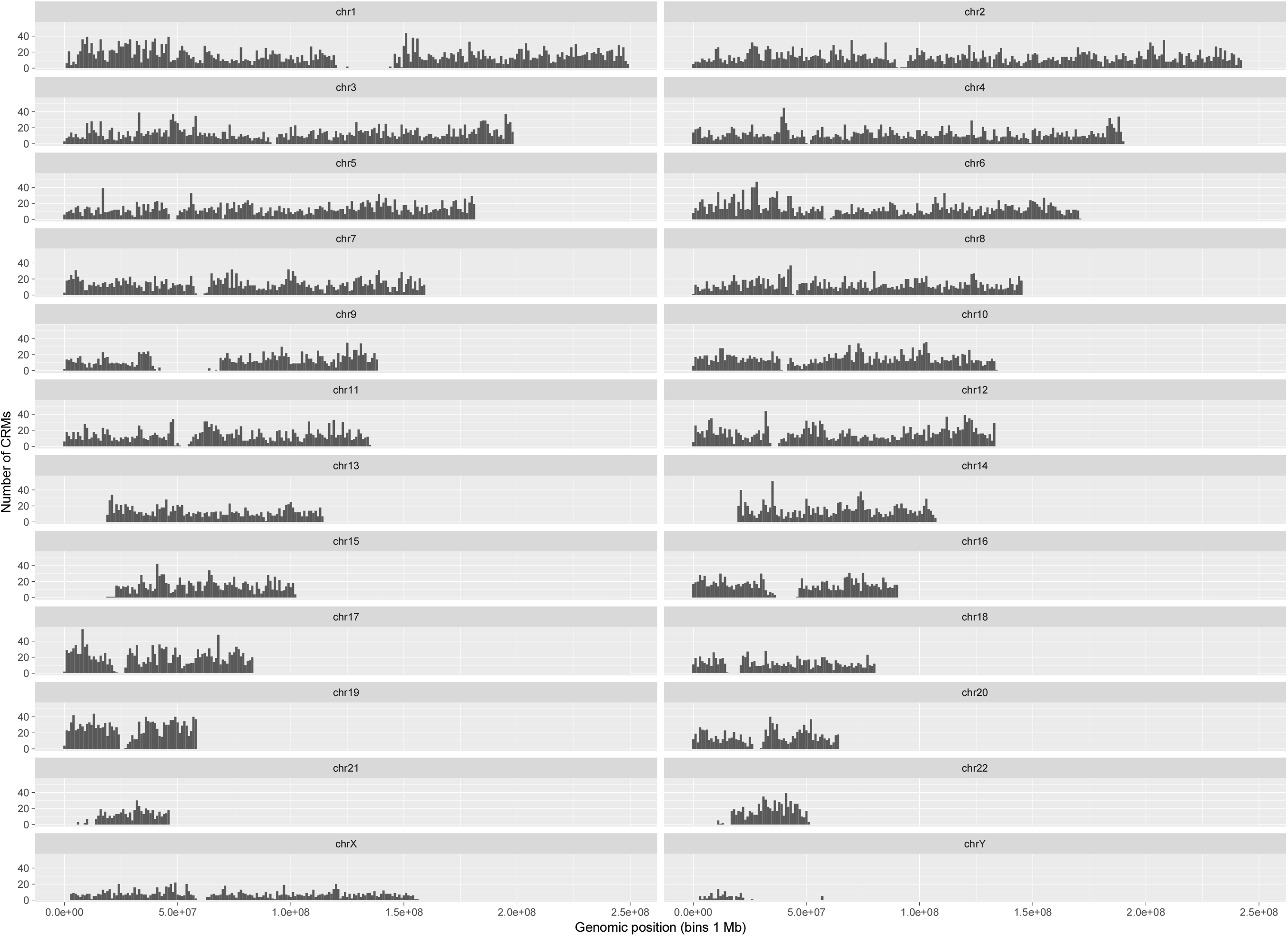
GRAMc CRM density over human genome 38, uniq.

**Supplementary Figure 5b.**
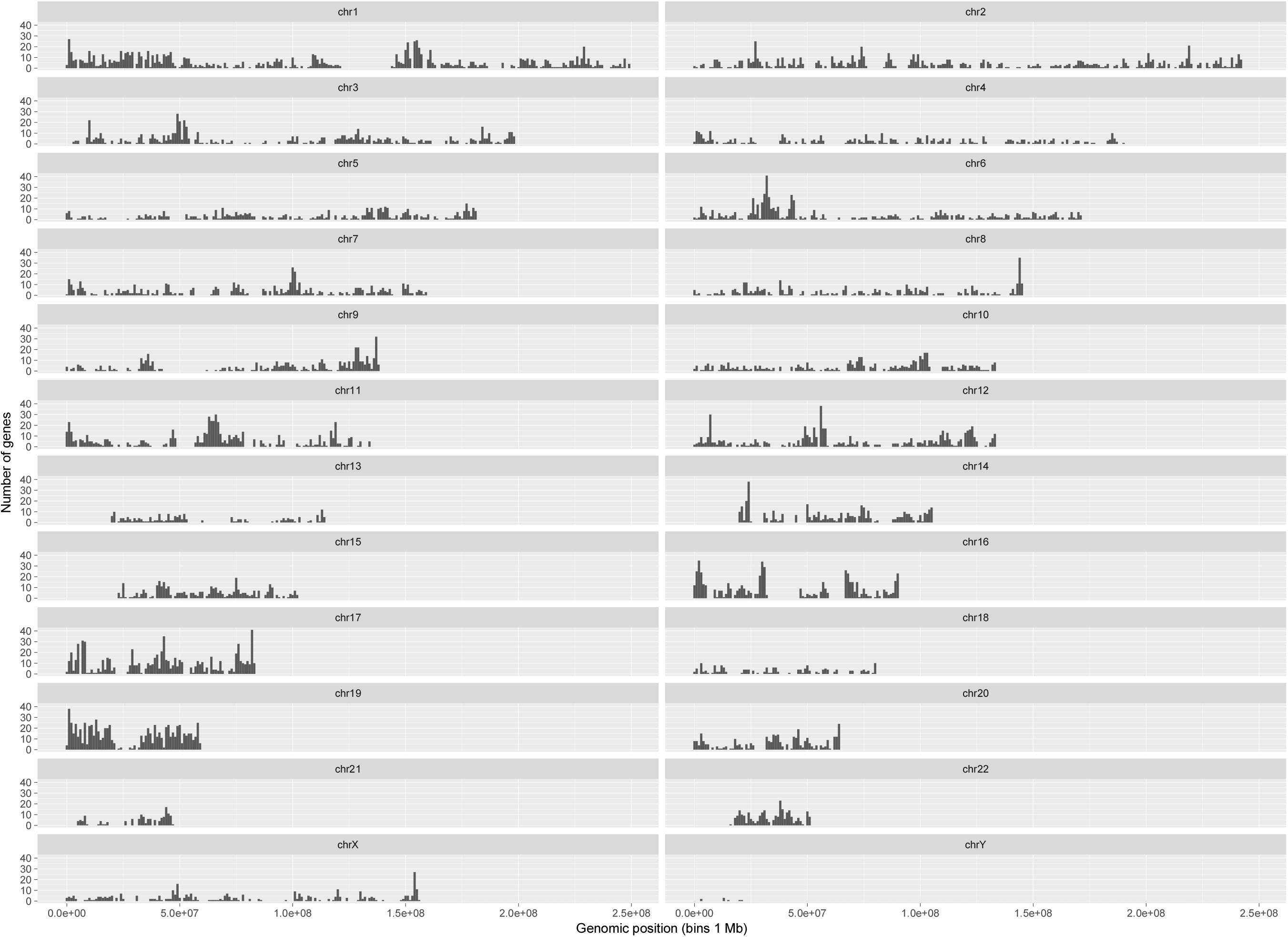
Expressed gene density over human genome 38.

**Supplementary Figure 5c.**
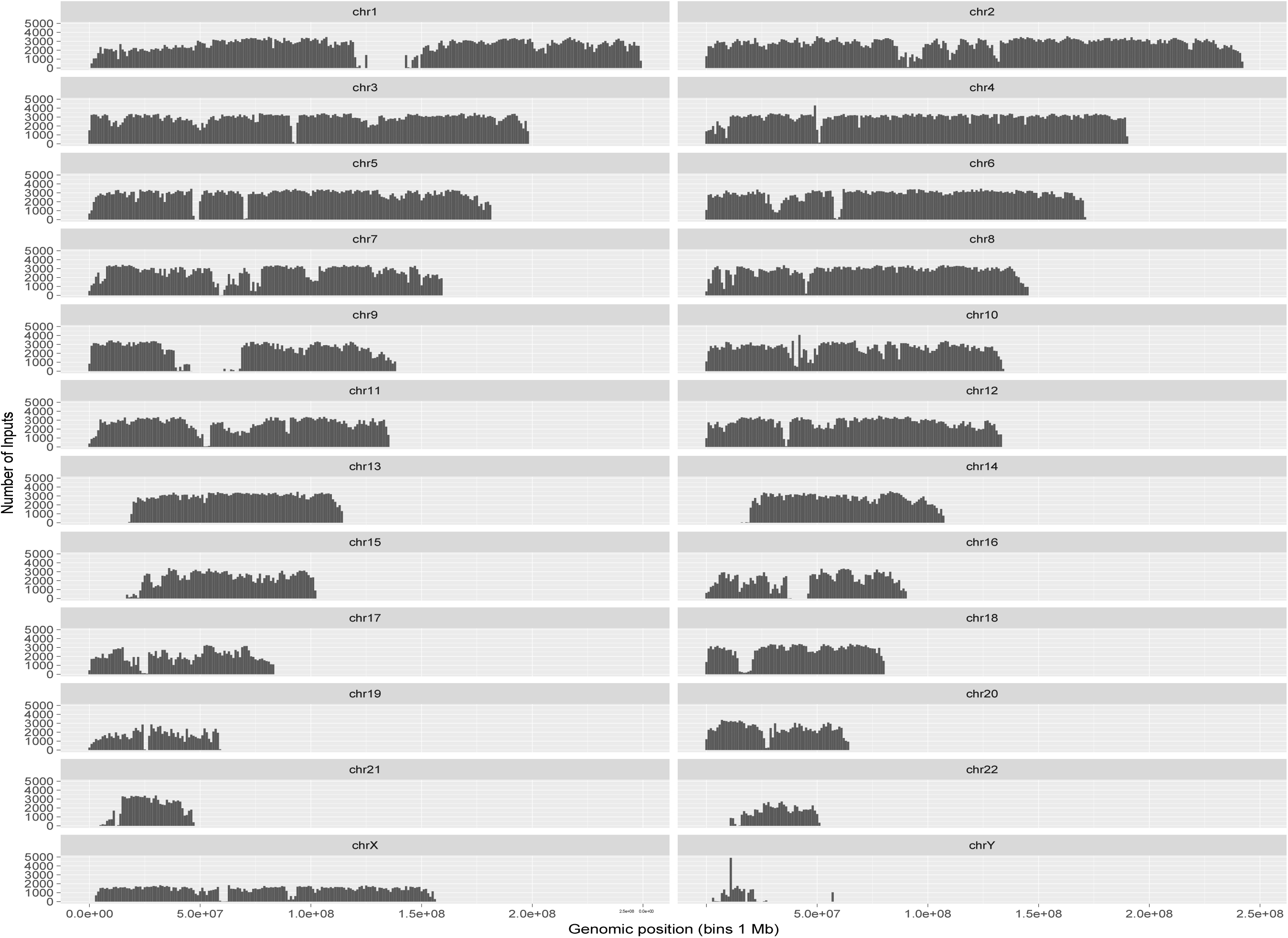
GRAMc Input density over human genome 38, uniq.

**Supplementary Figure 6.**
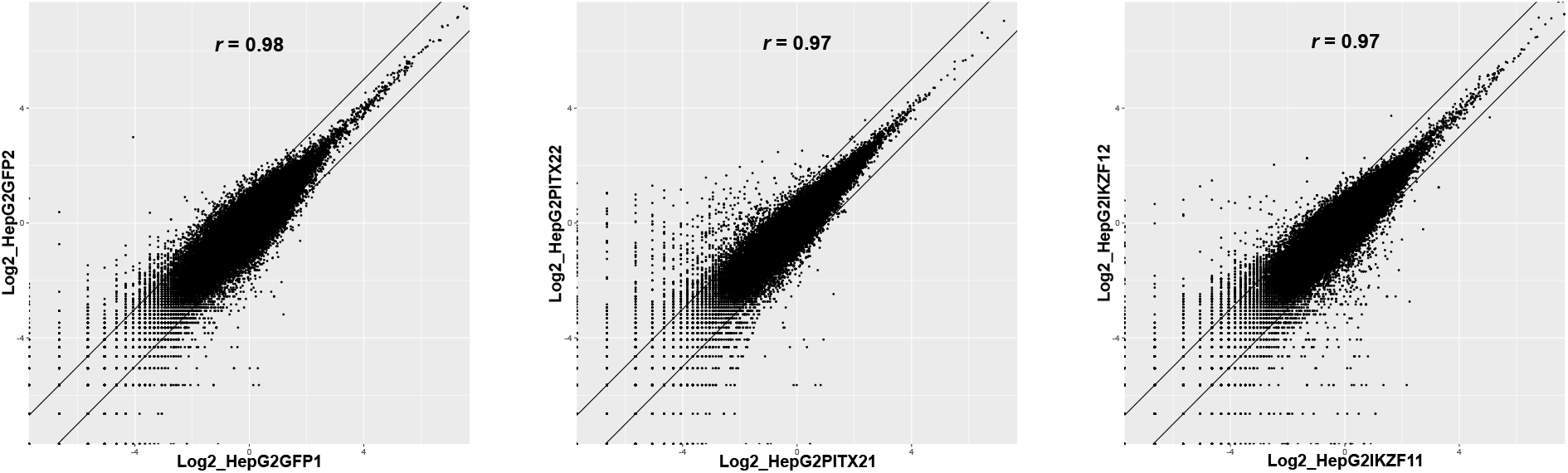
Reproducibility of perturbation experiments. Two independent batches of 80,000 randomly selected reporter constructs were compared for each perturbation experiment. All three experiments were highly reproducible (pearson’s r ≥0.97).

**Supplementary Files**:

https://rutgers.box.com/s/xctxdojtb3nre2hec8gg45kjqj8r1g5y

Supplementary File 1: List of GRAMc-tested inserts

Hs800PE_200MHepG2_B12_map50_bcl_5010_G0_hg38.uniq.bed

Supplementary File 2: List of ENCODE data files

SupplementaryFile.2.xlsx

Supplementary File 3: List of sequences used for motif enrichment analysis

SupplementaryFile.3.txt

Supplementary File 4: Motif enrichment data

SupplementaryFile.4.xlsx

Supplementary File 5: Enrichment of repeat families

SupplementaryFile.5.xlsx

Supplementary File 6: Primer sequences

SupplementaryFile.6.xlsx

Additional data files are available at https://rutgers.box.com/s/xctxdojtb3nre2hec8gg45kjqj8r1g5y

1. Sequence of template vector: (for Serial Cloner, http://serialbasics.free.fr/Serial_Cloner.html) SCP1_AflII-HindIII_Template_Plasmid.xdna

2. List of mapped inserts (hg38): Hs800PE_gDNA_All_map_mid.noalt.out

3. List of mapped inserts after removing non-unique inserts (hg38): Hs800PE_gDNA_All_map_mid.uniq.noalt.out

4. List of GRAMc-tested inserts before removal of non-unique sequences: Hs800PE_200MHepG2_B12_map50_bcl_5010_G0_hg38.bed

Note: Upon publication of the manuscript, all data files will be deposited on RUcore (Rutgers University Community Repository) for open access.

